# A foundation AI model enhances electron microscopy image analysis

**DOI:** 10.64898/2026.02.28.708664

**Authors:** Mengze Du, Yuwei Wang, Li Xie, Gaoliang Deng, Jiansheng Guo, Bo Han, Zhong-Hua Chen, Chen Rui, Jiankang Han, Yuan Chen, Yanru Zhao, Runzhou Cao, Fei Wang, Kun Li, Yu Wang, Yong He, Xuping Feng

**Affiliations:** College of Biosystems Engineering and Food Science, Zhejiang University, Hangzhou, China; College of Engineering, Anhui Agricultural University, Hefei, China; Analysis Center of Agrobiology and Environmental Sciences, Zhejiang University, Hangzhou, China; Hangzhou Chengfengerlai Digital Technology Co., Ltd, Hangzhou, China; Center of Cryo-Electron Microscopy, Zhejiang University School of Medicine, Hangzhou, China; Department of Computer Science, Hong Kong Baptist University, Hong Kong, China; School of Agriculture, Food & Wine, The University of Adelaide, Adelaide, Australia; College of Mechanical and Electronic Engineering, Northwest A&F University, Yangling, China; School of Computer Science and Information Engineering, Hefei University of Technology, Hefei, China; ReLER lab, CCAI, Zhejiang University, Hangzhou, China; The Rural Development Academy &Agricultural Experiment Station, Zhejiang University, Hangzhou, China

**Keywords:** electron microscopy, foundation model, image enhancement, mitochondrial ultrastructure, deep learning

## Abstract

Electron microscopy (EM) and advanced volume EM have extensive applications in deciphering cellular ultrastructures for life sciences. However, quality issues of EM images significantly impede the precise analysis and discovery of nanoscale biological structures. Here, we present an unsupervised Foundation Model for Five Tasks (DF5T) model for image enhancement through denoising, deblurring, super-resolution, two-dimensional (2D) inpainting, and three-dimensional (3D) isotropic restoration. DF5T was trained on an extensive dataset of multi-source membrane-bound organelle EM images, consisting of a total of over 2.25 million images. On unseen data, DF5T outperformed the existing state-of-the-art models in all five tasks. Furthermore, DF5T substantially compensates for missing three-dimensional structural information and reveals significant ultrastructural alterations in organelle geometry following chemical treatment. We demonstrate that DF5T significantly enhances electron microscopy images quality by improving the accuracy of downstream organelle segmentation and 3D structural restoration, thereby contributing to future advances in biological research.

## Introduction

The architecture and organization of organelle membranes, particularly the establishment of inter-organelle contact sites and junctions, are essential for numerous cellular processes^[1]^. Electron microscopy (EM), with its unparalleled nanoscale resolution, remains an indispensable tool for visualizing these structures and elucidating their functional significance^[2]^. Recently, three-dimensional (3D) EM techniques such as volume electron microscopy (vEM) and electron tomography (ET) have enabled 3D reconstruction of cells and tissues^[2]^. 3D EM has facilitated the discovery of novel structures^[3–6]^, enabled unprecedented insights into connectomics research, and has been recognized as one of the most promising emerging technologies in the life sciences^[7]^.

However, the power of EM is often compromised by inherent artifacts that compromise data fidelity. Pervasive issues—high noise levels, low contrast, and blur—severely hinder the accuracy of subsequent image segmentation and quantitative analysis^[7]^. This challenge is particularly pronounced in cryo-EM data from intact biological samples, where membrane structures often exhibit inherently low contrast against the cellular milieu, making their precise delineation arduous. Moreover, the serial nature of vEM leads to z-axis anisotropy, where resolution is lower between slices than within them. This complicates feature alignment across sections and leads to discontinuous or distorted 3D reconstructions of delicate structures such as the double membranes and mitochondria cristae, thereby compromising structural integrity^[8,9]^.

Although powerful, predominant deep learning strategies for EM restoration—such as PRS-SIM^[10]^, DBlink^[11]^, and DPA-TISR^[12]^ —rely on supervised learning, requiring large, curated datasets of paired low- and high-quality images. This requirement makes data acquisition and annotation costly and time-consuming. Additionally, these methods often use separate models for distinct tasks (e.g., denoising, super-resolution), which is inefficient for complex EM data degradation and can accumulate errors in sequential pipelines. They also require task-specific fine-tuning, limiting generalizability^[13–15]^.

A promising approach to addressing these limitations is the development of foundation models: large-scale models pre-trained on extensive datasets to learn generalizable feature representations. Such models can reduce the dependence on task-specific labels and be adapted to diverse downstream tasks^[16–18]^. By leveraging self- or weakly-supervised learning, these models mitigate the dependence on labeled data and can be adapted to various downstream tasks. Currently, the majority of foundation models are designed for fluorescence microscopy restoration^[17–19]^. However, applying this paradigm to EM presents unique and formidable obstacles. The physics of image formation in fluorescence microscopy and electron microscopy (EM) are fundamentally different, meaning a model trained on one may not transfer effectively to the other. Also, foundation models are notoriously data-hungry, and their performance scales with the size and quality of the training data. Therefore, high-quality, large-scale EM datasets are needed for foundation models, however, such datasets are exceptionally scarce and costly to acquire.

To address these challenges, we developed DF5T, an unsupervised, multitask, diffusion-based foundation model for EM image restoration. DF5T was trained on the largest collection of membrane-bound organelle EM images to date, comprising a total of over 2.25 million images. The model architecture uniquely leverages posterior sampling from a diffusion process, combined with singular value decomposition (SVD), to create a unified degradation framework. This allows DF5T to simultaneously perform denoising, deblurring, super-resolution, 2D inpainting, and 3D isotropic restoration within a single, coherent model. To validate its performance, we rigorously benchmarked DF5T against multiple state-of-the-art (SOTA) methods and demonstrate its superiority across all five tasks on unseen data. Critically, we show that the enhanced image quality from DF5T directly translates to significant improvements in downstream biological analysis, such as 2D organelle segmentation and 3D structural reconstruction. Finally, DF5T exhibits remarkable generalization, performing robustly across diverse EM modalities while showing adaptable potential for X-ray.

## Results

### Development and characterization of DF5T

Membrane-bound organelles define the compartmentalized architecture of eukaryotic cells, yet comprehensively resolving their intricate 3D topologies in electron microscopy (EM) images remains a major challenge. To address this, we introduce MemEM, a large-scale electron microscopy dataset specifically curated for membrane-bound organelles. MemEM integrates 1,111,771 real organelle membrane images derived from over 30 biological samples, encompassing diverse membrane structure types and functional states. Furthermore, 1,105,229 high-resolution, large-scale slice images were incorporated to better capture inter-organelle spatial interactions and three-dimensional context (Fig. 1a). To improve representation of long-tail and rare morphological features, we performed data augmentation on these minority classes using a text-guided Stable Diffusion model (SDM)^[20]^ (Supplementary Table 1 and Fig. 1b). By adopting the image-to-image paradigm with the original low-resolution electron microscopy images as strong guidance, we precisely controlled the generation process to produce highly realistic images while effectively preventing the introduction of artificial or false structural features. Experimental results demonstrate that models trained on MemEM significantly outperform baselines trained solely on real images across multiple super-resolution tasks, validating the dataset’s robustness and generalization capability in advancing systematic membrane structure analysis (Supplementary Fig. 1c-e).

**Fig. 1.**
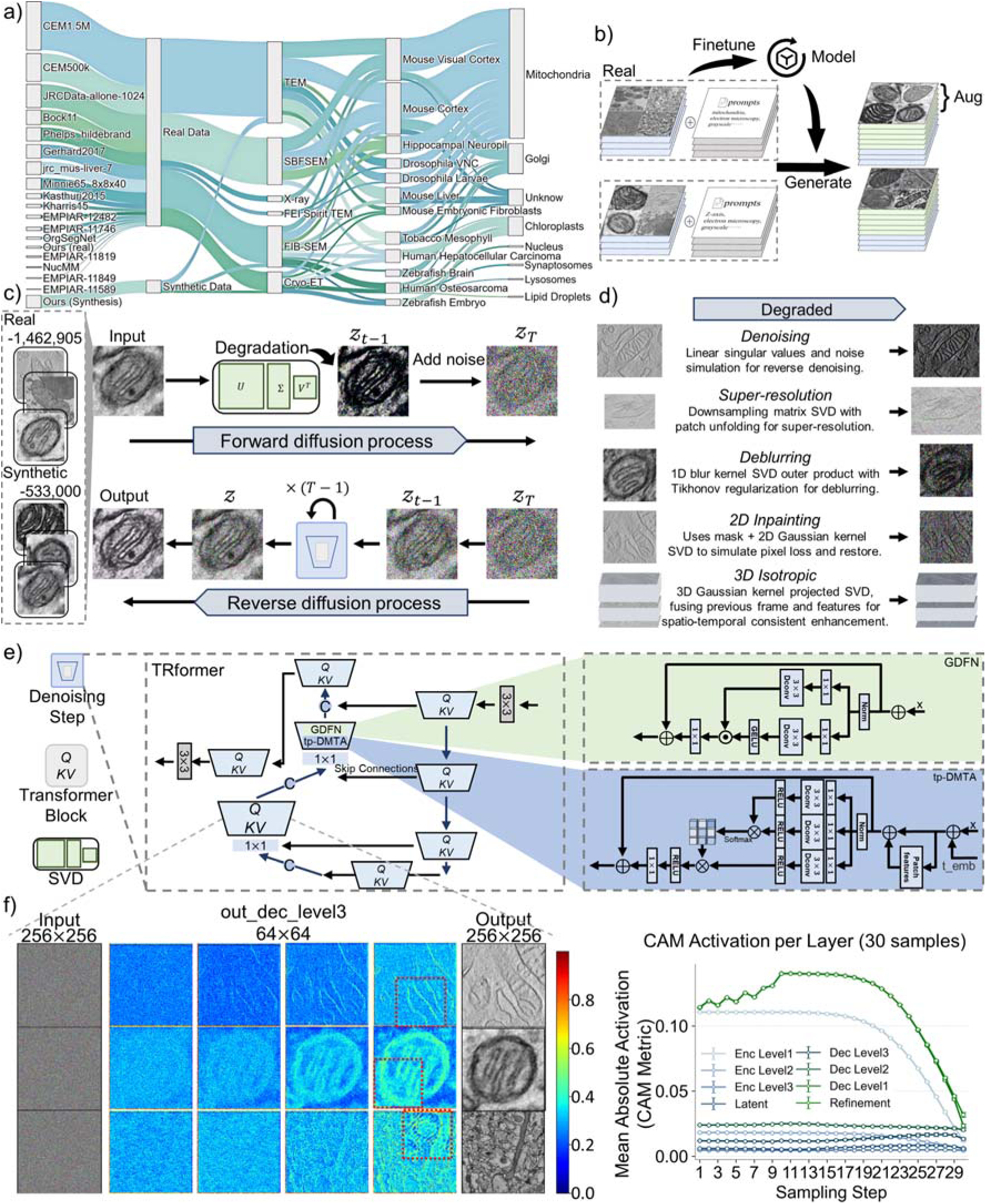
Review of multi-source data and model design strategies for the DF5T foundation model. a) Sankey diagram displays the distribution of dataset name, image type, imaging modality, species, and organelle. b) A data augmentation method based on synthetic images. c) Overall Architecture of DF5T. d) Unified degradation framework for multi-task micrograph enhancement. The model leverages singular value decomposition (SVD) to decompose the degradation process into five distinct tasks: denoising, super-resolution, deblurring, 2D inpainting, and 3D isotropic restoration. e) Transformer-based reverse denoising module. Each block within the module integrates a Gated-Dconv Feed-Forward Network (GDFN) and a Time-Patch Multi-Dconv Head Transposed Attention (tp-MDTA) mechanism, employing ReLU and GELU activation functions, respectively. f) Class activation map visualization. Class activation mapping of the out_dec_level3 layer, with bright regions (color intensity 0.6–0.8) indicating the model’s focus on areas within the image that align with our target membrane structures. Statistical distribution of CAM activation intensities across layers during the diffusion sampling process. For each layer, the mean absolute value was computed across all feature maps and all pixels in the layer’s output, representing the average activation intensity of the layer. Low fluctuation in this value indicates stable activation regions within the layer.

Based on the MemEM dataset, we developed DF5T (Fig. 1c)—an unsupervised model for image restoration. The model simulates diverse imaging modalities to progressively degrade images and reconstructs high-quality outputs through reverse diffusion, delivering comprehensive restoration capabilities including denoising, super-resolution, deblurring, 2D inpainting, and 3D isotropic reconstruction (Fig. 1d). To address the challenges of membrane structure characterization, we designed DF5T with a multi-scale encoder–decoder architecture that significantly enhances feature extraction of biological ultrastructure. A key innovation is the introduction of the temporal patch-wise multi-depth transposed attention (tp-MDTA) module, which maps temporal information into channel-specific features and integrates them with patch-wise representations extracted via adaptive pooling. This design effectively enhances the recognition of key subcellular structures. Complementing this, a gated deep feedforward network (GDFN) enables refined reconstruction of membrane continuity and morphological integrity (Fig. 1e). To elucidate the model’s interpretability, we visualized the reverse denoising process. DF5T initially reconstructs the global organelle morphology, then progressively refines membrane contours, and ultimately yields high-fidelity ultrastructural details in the final sampling steps. We further employed class activation mapping^[21]^ (CAM) to analyze the image restoration process and feature attribution patterns (Fig. 1f). Using the output from the “out_dec_level3” layer as input, we observed that high-activation regions (red) consistently localized to membrane structures. The mean absolute activation curves across 30 samples further demonstrated stable feature localization across network layers after only 30 sampling steps, confirming that our model has accurately captured organelle membrane characteristics from the MemEM dataset (Fig. 1g).

### Image Enhancement via Denoising, Deblurring, and Super-Resolution

In EM, low-dose imaging reduces electron counts, generating images with limited signal and mixed Poisson–Gaussian noise that degrades the signal-to-noise ratio (SNR). Lens aberrations, multiple scattering, and mechanical instabilities further blur high-frequency details and introduce misalignment, collectively limiting resolution. To address these multiple, partly orthogonal degradation modes, we decomposed image restoration into three specialized tasks—denoising, super-resolution and deblurring—and implemented each within DF5T via dedicated SVD-based modules^[22,23]^.

The denoising task addresses quantum noise–dominated degradation modeled as mixed Poisson–Gaussian noise, and uses an SVD-based operator to restore images. The super-resolution task enhances low-resolution images through SVD decomposition and inverse Gaussian kernel processing while maintaining spatial coherence. The deblurring task mitigates optical blur using a CTF-informed point spread function and a regularized SVD-based pseudo-inverse reconstruction with regularization for stability (Supplementary Figs. 3a, 6a and 9a). To evaluate the performance of our approach, we compared it with SOTA methods on a held-out validation set (Supplementary Table 2), quantifying image quality using peak signal-to-noise ratio (PSNR), structural similarity index (SSIM), and learned perceptual image patch similarity (LPIPS), and conducted comparative visualization experiments incorporating distinct biological validation methods across various tasks.

#### Denoising

A comparative analysis of image quality metrics across a range of resolutions demonstrates the feasibility and high-quality output of DF5T (Supplementary Fig. 3b). To validate its robustness, DF5T was applied to an unseen dataset, where it outperformed EMDiffuse^[23]^, Cellpose^[26]^, and UNiFMIR^[17]^ in denoising efficacy, as quantitatively confirmed by the higher frequency ratio (Fig. 2a). Critically, DF5T suppresses shot noise in cryo-ET slices of mouse embryonic fibroblast mitochondria while faithfully preserving ultrastructural membrane features—a capability quantitatively validated by membrane boundary gradient profiling (Fig. 2b). We also confirmed this robust performance across multiple additional datasets, including MemEM, Kasthuri2015, and Minnie65 (Supplementary Figs. 4–5), establishing DF5T’s broad applicability and generalization.

**Fig. 2.**
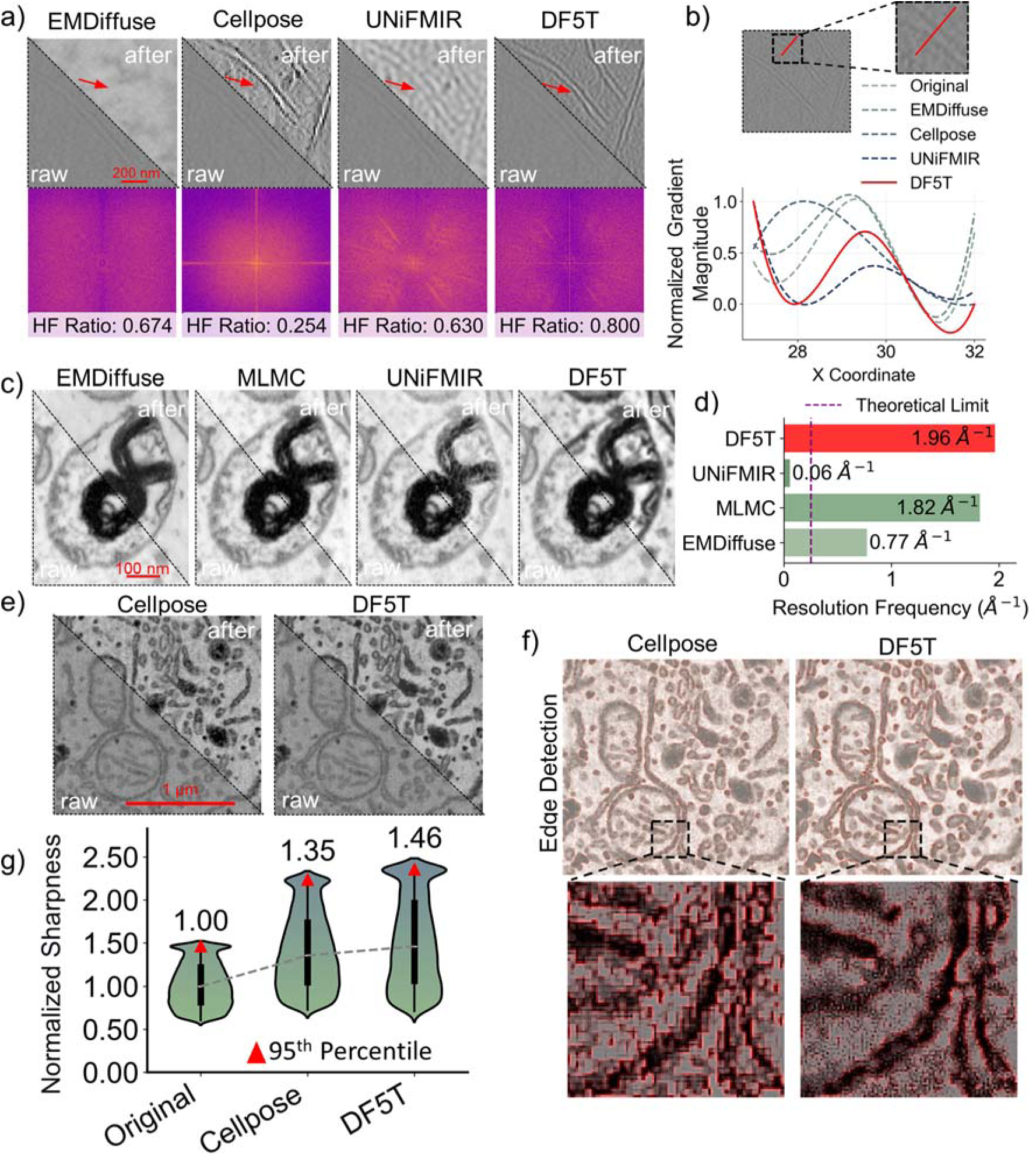
Performance of DF5T on denoising, super-resolution, and deblurring tasks. a) Denoising performance benchmarking on an unseen digital slice of cryo-electron tomography (cryo-ET) data from Mus musculus embryonic fibroblast mitochondria^[24]^. raw: input image. after: recovered images obtained by different algorithms. The HF ratio quantifies the relative contribution of high-frequency components in the image Fourier spectrum. A higher ratio indicates its ability to learn higher frequencies. b) Quantification of membrane boundary denoising through the normalized membrane boundary gradient profile. The peak values align with the original image, indicating robust preservation of the correct spatial positioning of membrane structures, while lower trough values signify superior denoising performance. c) Benchmarking super-resolution performance on one nucleus of focused ion beam scanning electron microscopy (FIB-SEM) dataset of a human bone osteosarcoma epithelial cell^[25]^. d) Quantification of super-resolution performance through Fourier Ring Correlation. The ‘Theoretical limit’ indicates the maximum resolution expected from the acquisition settings. A higher value indicates a better super-resolution effect. e) Benchmarking deblurring performance on one digital slice of FIB-SEM dataset of a human bone osteosarcoma epithelial cell^[25]^. f) Edge visualization for deblurring performance evaluation. A sobel filter is applied to compute the gradient magnitude, which is then normalized to highlight edge intensity. A semi-transparent red gradient layer, generated based on the normalized gradient magnitude, is overlaid on the contrast-enhanced image to accentuate regions exhibiting changes in edge sharpness. g) Quantification of membrane edge sharpness using sobel gradient magnitudes, normalized for cross-sample comparison.

#### Super-Resolution

DF5T effectively addresses the super-resolution problem (×2 upscaling) across a variety of data sources, outperforming EMDiffuse, UNiFMIR, and MLMC on key image quality metrics (Supplementary Fig. 6b). Visual inspection of super-resolution performance (Fig. 2c) further corroborates DF5T’s superior capability. Specifically, DF5T demonstrates superior reconstruction fidelity compared to other approaches, in accurately resolving intricate structural features such as densely packed membrane gap regions. This enhanced performance is evidenced by DF5T’s significantly reduced spatial distortion compared to MLMC and improved structural continuity relative to UNiFMIR, thereby establishing it as a more reliable solution for complex ultrastructural visualization. To quantify the spatial resolution of the reconstructions, we performed Fourier Ring Correlation to calculate the resolution frequency. Using a threshold of 0.143 (the 1/7 criterion), DF5T achieved a maximum spatial frequency of 1.96 Å ¹ (Fig. 2d). In four additional datasets, DF5T consistently produced the cleanest and most uniform structures in image reconstructions (Supplementary Figs. 7, 8).

#### Deblurring

DF5T demonstrated superior deblurring performance over advanced Cellpose, particularly in low-resolution imaging, where DF5T achieved median PSNR and SSIM values nearly three times higher (Supplementary Fig. 9b). Visual comparative analysis of deblurring results (Fig. 2e and Supplementary Fig. 10) further highlights key differences in representative images. While both methods improve clarity, DF5T resolves critical membrane features more effectively, an advantage highlighted by the edge-detection overlays. Magnified views revealed DF5T achieves a more tempered recovery of fine details, enhancing edge clarity without excessive prominence (Fig. 2f). This can be attributed to the EMDeblurring degradation module, which uses a CTF-informed point spread function to approximate the frequency response of the microscope optics, combined with a regularized SVD-based pseudo-inverse. This design captures the oscillatory contrast transfer and stabilizes the deconvolution at frequencies near CTF zeros, leading to sharper yet physically plausible membrane edges Quantitative analysis of normalized edge sharpness (see Methods for details) further revealed that DF5T achieves an optimal median sharpness value (Fig. 2g), surpassing Cellpose across all tested images (Supplementary Fig. 11).

### Image Restoration for 2D Inpainting and 3D Isotropic Restoration

During electron microscopy imaging, heterogeneous protein distribution across membrane structures produces elevated grayscale intensity in protein-enriched regions and reduced intensity in protein-sparse areas. Furthermore, sample preparation involving heavy-metal fixation and staining can introduce subtle structural compromises, resulting in discontinuities or aberrant connections within membrane architectures. In three dimensions, the distinctive lipid bilayer conformations, intermembrane spaces, and luminal domains of membrane-bound organelles form an intricate nanoscale network of membrane systems. However, pronounced anisotropy in 3D EM data leads to structural ambiguity, volumetric distortion, and challenges in serial image alignment (Supplementary Figs. 12a, and 14a).

In 2D inpainting, DF5T demonstrated superior performance and feasibility compared to the conventional restoration SDM model. Representative restored images revealed that DF5T outperformed SDM, particularly in inferring fracture points across different sample preparation methods, yielding enhanced structural detail and a more intact and continuous ultrastructure. This was quantitatively validated by metrics assessing sharpness, contrast, integrity, and edge characteristics (Fig. 3a, Supplementary Fig. 13).

**Fig. 3.**
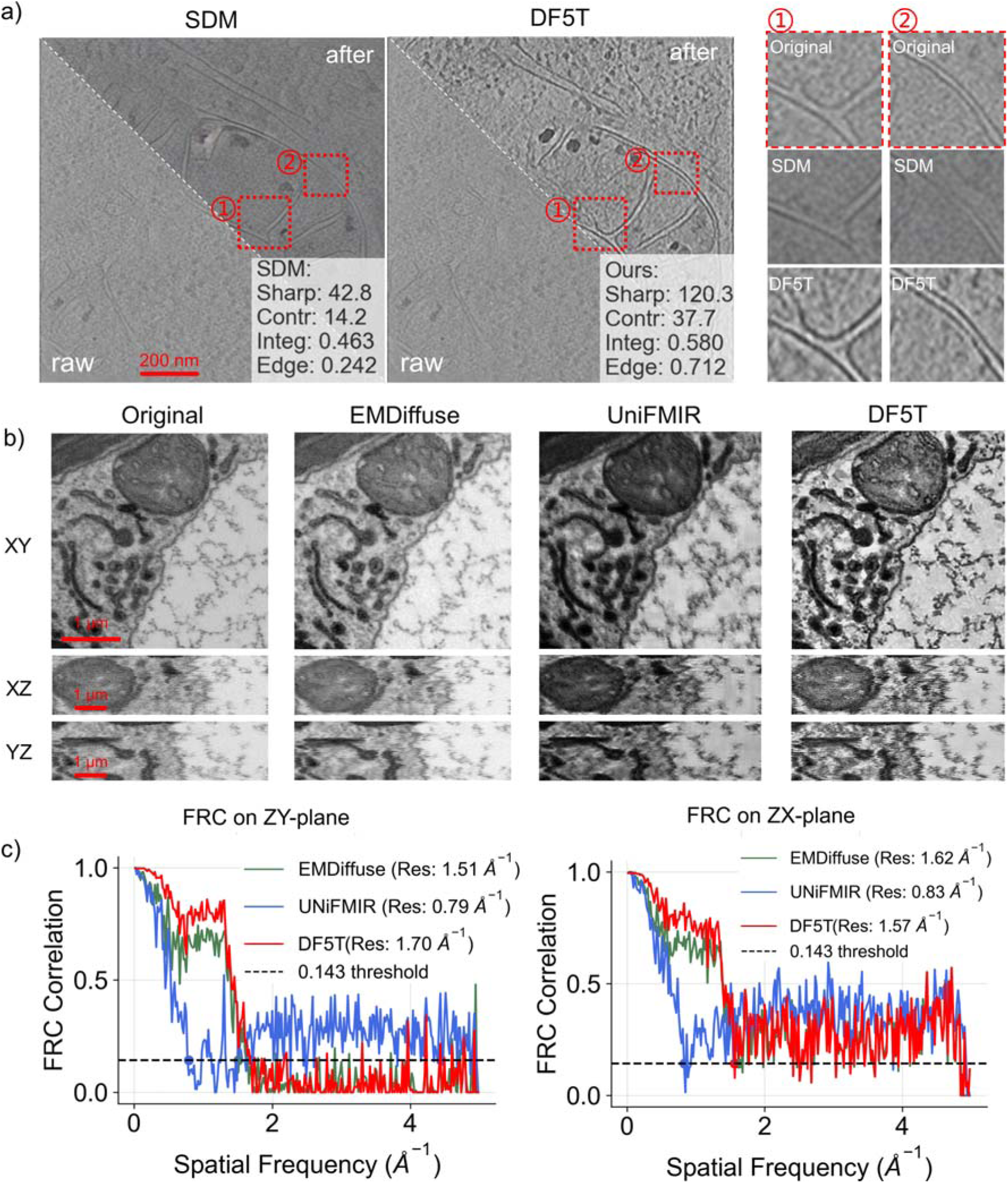
Performance of DF5T in 2D inpaint and 3D isotropic restoration tasks. a) 2D inpainting results on a representative cryo-ET slice of mitochondria from Mus musculus embryonic fibroblast^[24]^. Performance is quantitatively assessed by membrane sharpness, organelle contrast, structural integrity, and edge preservation, where superior scores denote enhanced image quality. Detailed views (① and ②) emphasize the accurate completion of partially damaged membrane structures. b) 3D isotropic restoration of mitochondria from tobacco leaf mesophyll cells^[5]^ (3D FIB-SEM). Orthogonal central slices through normalized 3D volumes (XY, XZ, YZ planes) illustrate the methodological improvements in capturing volumetric morphology and fine details. c) FRC resolution limits reveal superior z-axis anisotropy correction by DF5T. Resolution of YZ and ZX planes was assessed using Fourier Ring Correlation (FRC) metrics. Higher FRC resolution limits (Å□¹) at the 0.143 threshold on these planes directly indicate improved z-axis anisotropy correction and superior reconstruction fidelity.

For 3D isotropic restoration, we first constructed matrices from one-dimensional Gaussian kernels along the X, Y, and Z axes. Their subsequent SVD enabled the construction of separable convolution operations. This approach, combined with feature fusion using a pre-trained VGG model for perceptual similarity and adaptive weighting based on structural similarity and feature cosine similarity, alleviates axial–lateral resolution discrepancies in 3D EM data and yields more visually isotropic 3D reconstructions (Supplementary Fig. 14a). For cryo-ET, DF5T applies a 2D operator independently to each slice and does not use cross-slice information, thereby avoiding data leakage along the tilt axis. The method improves apparent smoothness and continuity of existing membranes. We compared images processed by DF5T with those restored using EMDiffuse and UNiFMIR, evaluating their performance through image quality metrics, 3D visualization, and isotropy analysis based on 3D fast Fourier transform (Fig. 4c-d, Supplementary Fig. 14b). DF5T consistently outperformed both methods across multiple image types (Fig. 3b-c, and Supplementary Fig. 15). Notably, DF5T’s 3D isotropic restoration demonstrated superior clarity and contrast, particularly in the ZY- and ZX-planes, as validated by both visual inspection and FRC metrics.

**Fig. 4.**
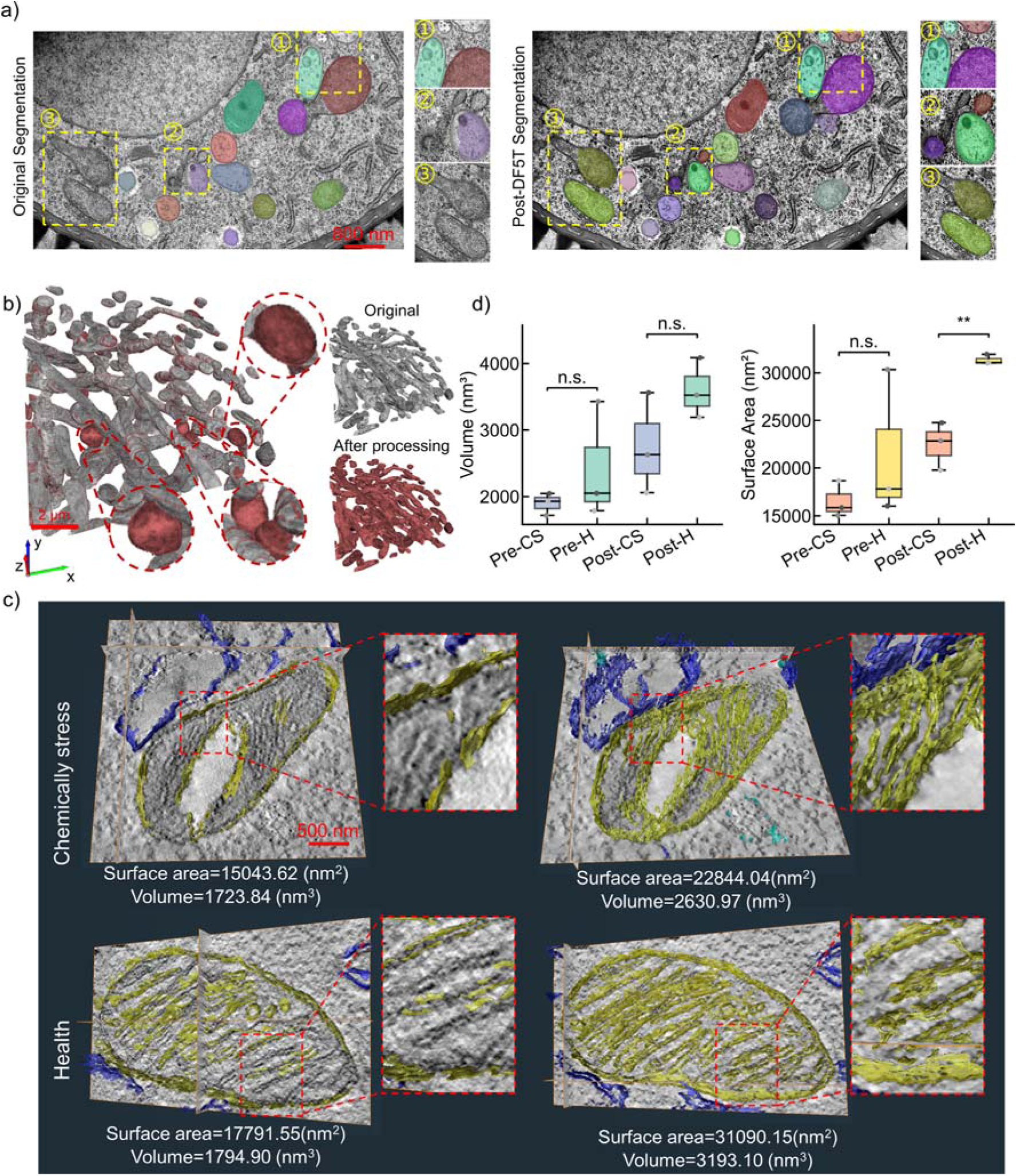
Enhanced segmentation and 3D reconstruction of cellular ultrastructure using DF5T. a) High-resolution transmission electron microscopy images^[27]^ were segmented using the Cellpose cyto3 model^[26]^ with the diameter parameter set to 300 pixels. Segmentation performance was compared between images pre-processed with Cellpose 3’s built-in denoising module and those pre-processed with the DF5T denoising module. b) 3D reconstruction of hippocampal neuropil from serial FIB-SEM sections^[28]^. The grey representation corresponds to data without DF5T processing, and the red representation shows the results after DF5T processing. The overlapping regions are prioritized in gray. We employed the 3D instance segmentation outputs produced by empanada. c) Mitochondrial ultrastructural analysis in chemically stressed hepatocytes. Electron tomography reconstructions demonstrate enhanced 3D segmentation fidelity after DF5T processing (right) compared to baseline methods (left), specifically under oleic acid treatment conditions. d) Quantitative comparison of mitochondrial morphology under chemically stressed (CS) versus healthy (H) conditions (ANOVA). Boxplots display volume (left) and surface area (right) measurements pre- and post-DF5T processing (n=3). **. p<0.01; n.s. p≥0.05.

### External evaluation in downstream tasks after DF5T processing

The quality of raw EM data critically influences downstream image analysis. Therefore, we evaluated the effect of DF5T processing on the 2D segmentation and 3D reconstruction of EM datasets (Fig. 4 and Supplementary Figs. 16-17). Following DF5T application, we observed significantly enhanced clarity of organelle boundaries and effective noise suppression. This resulted in a higher average Intersection over Union (IoU) for organelle segmentation errors (Fig. 4a).

To further validate the power of DF5T for 3D reconstruction, we examined published mouse cortex vEM data from serial FIB-SEM sections^[29]^. Our analysis revealed that DF5T processing renders more complete and distinct mitochondrial membrane boundaries. This improvement led to an increased number of identified mitochondrial regions and a significant decrease in falsely detected connected regions (Fig. 4b). For ET data from another cultured mouse hepatocyte cell lines, our results show that unprocessed ET images exhibit substantial noise, making it challenging to effectively segment individual mitochondria. Blurred membrane boundaries result in non-smooth mitochondrial morphology in 3D reconstructions, with missing segments of cristae or the presence of numerous voids. DF5T processing of both healthy and oleic acid-treated samples led to increased mitochondrial volume and surface area with enhanced membrane boundary definition, improved cristae visibility, and significantly reduced false topological connections. Specifically, the difference of mitochondrial surface area increased from 15% to 27%, and the difference in volume grew from 4% to 18% (Fig. 4c and d). Crucially, DF5T processing preliminarily revealed a statistically significant difference (p<0.01) in mitochondrial surface area between the two treatment groups, a key biological distinction that was completely masked in the unprocessed images. These findings provide critical evidence for understanding the impact of chemical stress on sample characteristics.

### DF5T generalizes across serial processings and imaging modalities

The correspondence between color noise patterns and the enrichment of high-frequency dark regions with membrane edge characteristics aligns with their intrinsic role as high-contrast, high-frequency components. Complementary class activation mapping overlays reveal strong spatial correspondence between high-activation regions and membrane boundaries in high-frequency images, confirming that DF5T restores membrane structures by leveraging authentic high-frequency features (Fig. 5a). Moreover, the consistent alignment of these high-frequency features throughout the restoration process highlights DF5T’s robust generalization capability, enabling effective performance across a range of downstream tasks (Fig. 5b). This ability to preserve high-frequency information ensures stable performance even across multiple sequential processing stages.

**Fig. 5.**
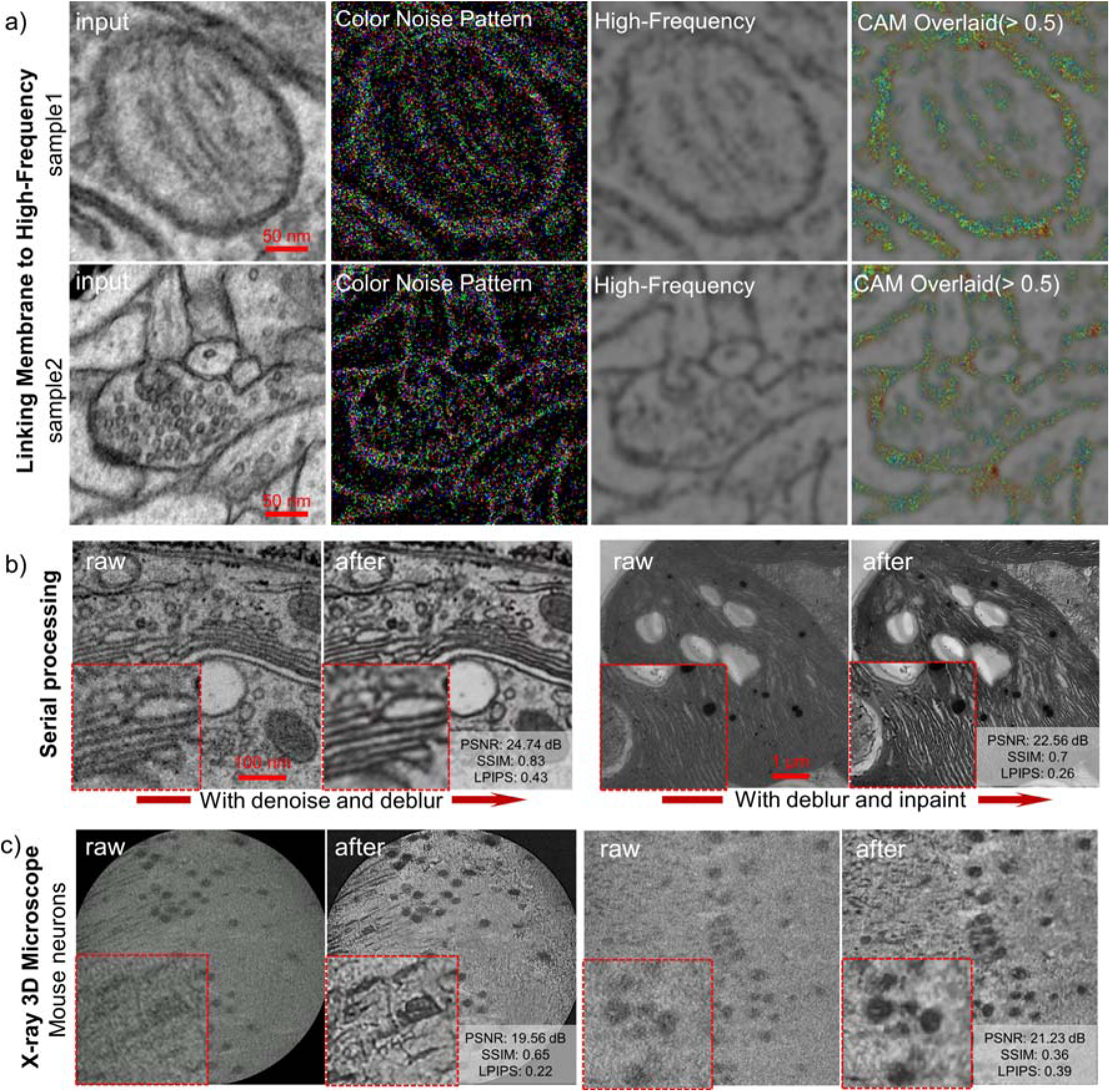
Generalization of DF5T across organelles and imaging modalities. a) Correlation Between Membrane Features and High-Frequency Information. Color Noise Pattern shows the out_dec_level3 activation map at 30-step process. High-Frequency utilizes high-pass filtering (subtracting low-frequency components after Gaussian blur) to separate high-frequency components from the input image. The CAM overlaid image superimposes the regions with activation intensity values greater than 0.3 in the activation map onto the high - frequency image. b) Comparison of different approaches for addressing imaging challenges, including sequential pipelines involving stacked tasks. c) In a zero-shot capability, DF5T applied to 3D X-ray microscopy data of mouse neurons, showing improved structural clarity and detail.

In a zero-shot paradigm, this enhancement of high-frequency features also extends to other imaging modalities; for example, DF5T produces sharper structural details in 3D X-ray microscopy of mouse neurons (Fig. 5c). Quantitative metrics, including improvements in PSNR, SSIM, and LPIPS, further validated these gains, characterized by enhanced structural clarity and reduced noise.

## Discussion

We present DF5T, an unsupervised diffusion-based foundation model designed to address five critical EM image degradation challenges: denoising, super-resolution, deblurring, 2D inpainting, and 3D isotropic restoration. DF5T is applicable across diverse EM modalities, including 2D SEM and TEM as well as 3D vEM and ET. Across all restoration tasks and modalities, DF5T consistently outperformed existing SOTA methods, demonstrating robust generalization on unseen datasets (Fig. 5).

DF5T substantially improved efficiency in EM image analysis. In 2D EM, improved image quality facilitates organelle segmentation with greater accuracy. In 3D EM, DF5T resolves intricate membrane structural features, such as membrane contact sites and organelle membrane remodeling^[30]^, which are essential for understanding inter-membrane material exchange and organelle function. By mitigating z-axis anisotropy and membrane discontinuities, DF5T enhances high-frequency membrane features along the z-axis, yielding smoother and more complete 3D reconstructions with improved segmentation accuracy, clearly delineating membrane boundaries and continuous organelle morphology.

Unlike supervised models (e.g. EMDiffuse and DBlink) or fluorescence-specific models (e.g. UNiFMIR and SACD^[31]^), DF5T operates as an unsupervised foundation model trained on approximately 2.25 million EM images. Its SVD-based degradation modeling, tailored to EM data characteristics, adaptively suppresses noise using singular values and anisotropy parameters, stabilizes deblurring via regularized pseudoinverse operations, simplifies super-resolution through block-wise mapping, guides 2D inpainting with joint edge detection, and addresses 3D anisotropy using anisotropic Gaussian–based operators and, for vEM data, optional cross-slice contextual information (Fig. 1). Integrated with the Restormer architecture, featuring tp-MDTA and GDFN, DF5T precisely restores low-to-high-resolution images by learning structural features. Class activation map visualizations confirm progressive focus on key organelle structures, aligning with human recognition patterns.

DF5T demonstrates a remarkable ability to process high-frequency features in EM data, highly beneficial for discerning organelle membrane structures. Crucially, DF5T demonstrates exceptional generalization capabilities, marking a groundbreaking advancement in EM image quality enhancement across diverse organelle membrane structures, and DF5T achieves remarkably superior performance in regions of interest across other imaging modalities without requiring fine-tuning. While DF5T demonstrates robust generalization across diverse membrane structures and imaging modalities, its computational demands—particularly its reliance on large-scale datasets and high resource requirements—remain a practical limitation. Future optimizations, such as few-shot learning, lightweight architectures, or token-based reconstruction objectives^[32]^, could significantly reduce these constraints. Such advancements would enhance DF5T’s accessibility and solidify its role as a pivotal tool for ultrastructural and systems biology research.

## Methods

### MemEM dataset acquisition and pre-processing

To support the training of foundation models with both high quality and strong generalization capabilities, we constructed the MemEM dataset, a large-scale collection of membrane-bound organelle images comprising 2,250,000 images (see Supplementary Table 1). The real data were primarily sourced from 17 public datasets, supplemented by newly acquired electron tomography (ET) datasets derived from mouse myoblast cell lines and tobacco leaves. To highlight the breadth of MemEM, it encompasses diverse species categories, including mammals (such as mouse and human), insects (such as Drosophila), nematodes (such as Caenorhabditis elegans), fish (such as zebrafish), and plants (such as tobacco and 19 other species), and incorporates multiple imaging modalities, including transmission electron microscopy, serial block-face scanning electron microscopy (SBEM), focused ion beam scanning electron microscopy (FIB-SEM), and cryo-electron tomography (cryo-ET). The 3D-EM datasets were sliced along the X, Y, and Z axes at their respective resolutions to generate 2D EM images. Segmentation of membrane-bound organelles was performed using the micro-sam, empanada, and OrgSegNet models, all with default parameters. Following exclusion of segmented regions smaller than 128 × 128 pixels and uninformative patches, this yielded approximately 2.25 million low-resolution membrane-bound organelle images.

### Electron tomography analysis

All the samples were fixed with 2.5% glutaraldehyde overnight, followed by 1% osmium tetroxide in 0.1 M phosphate buffer (pH 7.0) at room temperature (1.5 h for cell lines and animal tissues, 2 h for plants). Each fixation step was followed by three 15-minute rinses with 100 mM phosphate buffer (pH 7.0) to remove residual fixatives. Dehydration was performed using a graded ethanol series (50%, 70%, 80%, 90%, 95%, 100%) and pure acetone, each for 20 minutes at room temperature. Samples were embedded in Epon 812 resin (SPI Supplies Inc., PA, USA) and polymerized at 60°C for 48 hours to form solid blocks preserving ultrastructure. Ultra-thin sections (200 nm) were cut using a Leica UC7 microtome (Leica, Vienna, Austria) with a diamond knife (Diatome, Switzerland) and mounted on 100-mesh carbon-coated copper grids. Sections were double-stained with uranyl acetate and lead citrate for 15 minutes each, followed by application of 10 nm colloidal gold particles (Sigma, St. Louis, MO) for 5 minutes on both sides to facilitate ET data alignment. Mitochondrial data were collected automatically using the JEOL Recorder program on a JEM-2100 Plus TEM (JEOL, Tokyo, Japan) at 200 kV, with images captured by a bottom-mounted Rio9 CMOS camera (3k × 3k, Gatan, CA, USA). Acquisition parameters included ×12,000 magnification, 2.83 e/sÅ² beam dose, 0.546 nm pixel size, tilting from −60° to +60° in 1° increments, 6 μm defocus, and binning=1. 3D reconstruction was performed using the IMOD eTomo program, employing fiducial model generation, alignment, tomogram positioning, and tomogram generation via back-projection and simultaneous iterative reconstruction (SIRT) algorithms34,35. Fine alignment used at least 30 gold particles per tilt series, achieving a residual error below 0.5^[33,34]^.

### Training dataset preprocessing

To address heterogeneity in formats and contrast, images underwent a three-step preprocessing pipeline (Supplementary Fig. 2a). First, an adaptive grayscale threshold (70–100%) extracted organelle membranes; second, a density threshold (35–100%) removed artifacts such as noise and clustered shadows, enhancing membrane details (Supplementary Fig. 2b); third, optical flow–based interpolation leveraged continuity across frames to restore missing membrane segments (Supplementary Fig. 2c). This produced standardized, high-quality images for training the foundation super-resolution model.

### Data augmentation method

Synthetic EM images were generated using a Stable Diffusion–based pipeline. To augment the multi-source dataset, ∼2 million images were annotated with species, 3D-EM axis orientation, data type, and image format, then used to fine-tune stable-diffusion-v1-5 into a low-rank adaptation of the Stable Diffusion model optimized for EM features (Fig. 1a). To enhance performance in data-scarce categories, we focused on generating data for four tail classes: cryo-electron tomography Golgi (EMPIAR-11589), mitochondria (EMPIAR-11819), and chloroplasts (EMPIAR-12482), along with a limited quantity of FIB-SEM chloroplast data (The Plantorgan hunter dataset). The generation process incorporated manually curated high-quality image prompts paired with text prompts derived from data type and category. This ultimately yielded 1,000 cryo-ET Golgi images and a total of 30,000 images comprising cryo-ET mitochondria, cryo-ET chloroplasts, and FIB-SEM chloroplasts through generative modeling. LoRA fine-tuning was performed using sd-trainer, with training images including mitochondria (128–512 px) and slice images (256–1024 px). Key training parameters were as follows: UNet learning rate of 1×10□□, text encoder learning rate of 1×10□□, network dimension of 32, and 10 training epochs. The final generative model was created by merging the fine-tuned LoRA adapter with the base Stable Diffusion WebUI model at a weight of 0.3. Image generation employed the Diffusion Probabilistic Model++^[35]^ with 30 sampling steps, producing 33,000 images at uniform resolutions. Model training utilized stochastic gradient descent with layer-wise adaptive rate scaling, momentum, and weight decay. Training was conducted on four NVIDIA H800 GPUs (effective batch size of 2) for 10 epochs, requiring approximately 5 days.

### Constructing a Diffusion-based Foundation model for 5 Tasks

The DF5T framework builds upon classical diffusion model architecture to enable efficient unsupervised posterior sampling. It enhances control over generated sample characteristics by incorporating additional conditional information, achieving greater precision and quality during the generation process. The reverse process of the DF5T model follows a parameterized Markov chain. In this reverse process, the denoising step is defined as:

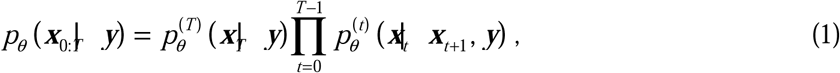

where *y* serves as the conditioning guide, and *x*_0_ represents the final diffusion output. The factorized variational distribution conditioned on *y* is implemented as:

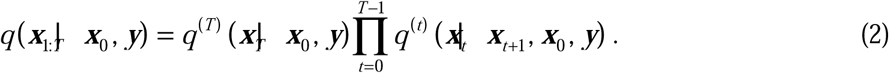

This establishes a variational lower bound objective for the diffusion model conditioned on *y*, enabling the denoising process to approximate the probability distribution of recovering noise-free observations under the condition *y*.

The DF5T architecture comprises two primary components: a multitask degradation method based on SVD (Fig. 2b) and a denoising module built on an enhanced Transformer (Fig. 2c).

Inspired by the denoising diffusion restoration model network^[36]^, the multitask degradation method integrates SVD into our framework, tailored to the unique degradation characteristics of EM data. It is differentiated into denoising, deblurring, super-resolution, 2D inpainting, and 3D isotropic restoration tasks (Fig. 2b).

The improved backpropagation module is illustrated in Fig. 2c. Based on the Restormer^[37]^ framework, three enhancements are introduced to address the characteristics of our dataset. First, unlike the classic UNet architecture, a TRformer denoising module replaces the original UNet denoising module. Within the attention mechanism, the tp-MDTA is designed. Through temporal embedding, combined with adaptive pooling and bilinear interpolation, partitioned spatio-temporal feature fusion is achieved. This approach integrates global contextual information with local details, effectively reducing the computational complexity of high-resolution features and improving the capture of complex membrane structures. It is particularly suited for handling highly repetitive texture patterns in EM images. Additionally, temporal embedding (Emb) projects the temporal conditions of the diffusion model into the feature space via channel expansion (expand), enabling dynamic weighted fusion of spatio-temporal features. A learnable temperature parameter *τ* dynamically adjusts the attention distribution, significantly improving the efficiency of high-resolution image processing, defined as:

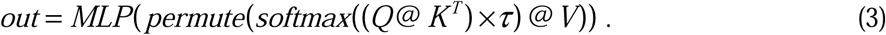

Temporal embedding is implemented as:

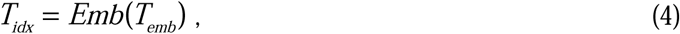

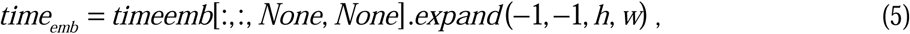

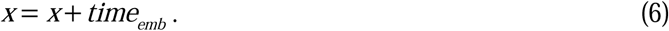

Partitioning is achieved through the following formulations:

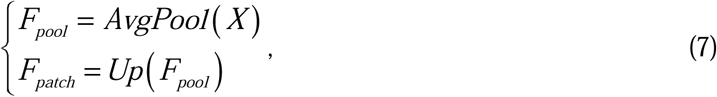

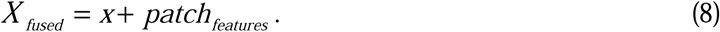

The resulting query (Q), key (K), and value (V) are obtained as:

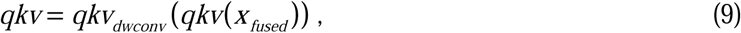

Second, a dynamic gating mechanism is introduced in the feed-forward network, employing a dual-path parallel convolution structure for feature interaction. Feature selection is achieved through GeLU activation and element-wise multiplication, enhancing nonlinear expressiveness compared to traditional feed-forward networks. Third, a hybrid downsampling module combines transposed convolution and standard convolution for dual-path downsampling, preserving high-frequency details while enabling multi-scale feature extraction, outperforming traditional UNet pooling operations. A four-level encoder-decoder architecture with cross-layer skip connections enables simultaneous capture of high-frequency details dominated by organelle membranes and low-frequency contours comprising imaging noise and cytoplasmic components. Overlapping convolution kernels further mitigate edge artifacts caused by traditional partitioning. Temporal embedding maps diffusion model time-step information to the feature space via linear projection, integrating with the key-value matrices in the attention mechanism. This allows the network to dynamically adjust feature weights based on noise intensity, achieving deep coupling of temporal and spatial conditions. Ablation study results are provided in Supplementary Table 3.

### Computational Resources for Base Model Training

To enable high-fidelity 3D generation of cellular organelles from electron microscopy (EM) data, we developed DF5T, a diffusion-based multi-modal large model trained on MemEM. DF5T was implemented in PyTorch, conditioned on EM technique and organelle class, and trained for 500,000 iterations on four NVIDIA H800 GPUs (effective batch size 2 via gradient accumulation, initial learning rate 5 × 10□□, AdamW optimizer with cosine annealing scheduler, and linear noise schedule from *β*_0_ =0.0001 to *β_T_* =0.02. Ablation studies (Fig. 4f) were performed using identical parameters on a subset of 50,000 images to evaluate the contributions of individual enhancements. Visualizations were generated using Matplotlib and Seaborn libraries in Python.

### Feature visualization and saliency maps

Feature visualization and saliency maps were generated using CAM techniques (Fig. 1d). Feature maps from the third decoder layer (‘out_dec_level3’) were analyzed to reveal the model’s spatial attention during image processing. CAM computation involved channel-wise averaging of feature maps to determine mean activation values at each spatial location, quantifying attention intensity across regions. Activation maps were subjected to min-max normalization, scaling values to the [0, 1] interval to standardize scales and enhance contrast. To improve biological interpretability, a multi-scale feature fusion strategy was incorporated, employing adaptive pooling and bilinear interpolation during forward propagation to refine capture of fine-grained structures. The resulting CAM outputs were rendered as high-resolution heatmaps using the Jet colormap, highlighting focus on key biological elements, such as cellular boundaries, textures, or pathological features.

### Graphical user interface design

A graphical user interface was developed to improve accessibility and user interaction. This interface supports processing of electron microscopy images, incorporating five core functions. Based on DF5T, the graphical user interface allows adjustment of time steps and noise levels to optimize image quality, thereby enhancing visibility of membranes and internal structures. Implemented in PyQt6, the interface provides real-time previews and result comparisons. A step-by-step operational demonstration is available in the supplementary video tutorial (https://www.youtube.com/watch?v=2AW3lW8pVhw).

## Data Availability

All data used in the experiments are accessible for browsing and download via the links provided in Supplementary Table 1. The MemEM multi-source dataset constructed for this study is available at https://www.scidb.cn/detail?dataSetId=8d30c6b23acd46d09e44114e8f739fe4#p3.

## Code Availability

The source codes of DF5T, several representative pre-trained models as well as some example images for testing are publicly accessible via https://github.com/dumengze/DF5T.git.

## Acknowledgements

The authors thank Professor KANG Byung Ho from the School of Life Sciences, The Chinese University of Hong Kong, for his insightful guidance and meticulous review, which greatly improved the quality of this paper.

## Author contributions

Data curation: M.D. Methodology: M.D., R.C., F.W., K.L., and Y.W. Resources: G.D., C.R., J.H., and Y.C. Funding acquisition: L.X., J.G., X.F., and Y.H. Visualization: M.D., X.F., and Y.W. Software: M.D. Supervision: B.H., Z.-H.C., and Y.Z. Validation: X.F., and M.D. Writing—original draft: X.F., M.D., and L.X. Writing—review and editing: X.F., and M.D.

## References

K. W. Ryu, T. S. Fung, D. C. Baker, M. Saoi, J. Park, C. A. Febres-Aldana, R. G. Aly, R. Cui, A. Sharma, Y. Fu, O. L. Jones, X. Cai, H. A. Pasolli, J. R. Cross, C. M. Rudin, C. B. Thompson, Nature 2024, 635, 746.

N. L. Turner, T. Macrina, J. A. Bae, R. Yang, A. M. Wilson, C. Schneider-Mizell, K. Lee, R. Lu, J. Wu, A. L. Bodor, A. A. Bleckert, D. Brittain, E. Froudarakis, S. Dorkenwald, F. Collman, N. Kemnitz, D. Ih, W. M. Silversmith, J. Zung, A. Zlateski, I. Tartavull, S. Yu, S. Popovych, S. Mu, W. Wong, C. S. Jordan, M. Castro, J. Buchanan, D. J. Bumbarger, M. Takeno, R. Torres, G. Mahalingam, L. Elabbady, Y. Li, E. Cobos, P. Zhou, S. Suckow, L. Becker, L. Paninski, F. Polleux, J. Reimer, A. S. Tolias, R. C. Reid, N. M. Da Costa, H. S. Seung, Cell 2022, 185, 1082.

X. Li, F. Qiao, J. Guo, T. Jiang, H. Lou, H. Li, G. Xie, H. Wu, W. Wang, R. Pei, S. Liu, M. Ye, J. Li, S. Huang, M. Zhang, C. Ma, Y. Huang, S. Xu, X. Li, X. Sun, J. Yu, K. L. Fok, S. Duan, H. Chen, Nat Commun 2025, 16, 1664.

G. Wang, J.-S. Guo, H.-J. Huang, Z.-R. Zhu, C.-X. Zhang, Commun Biol 2025, 8, 441.

J. Guo, G. Wang, L. Xie, X. Wang, L. Feng, W. Guo, X. Tao, B. M. Humbel, Z. Zhang, J. Hong, Plant Cell & Environment 2023, 46, 650.

X. Wang, J. Ma, X. Jin, N. Yue, P. Gao, K. K. K. Mai, X. Wang, D. Li, B. Kang, Y. Zhang, JIPB 2021, 63, 353.

C. J. Peddie, C. Genoud, A. Kreshuk, K. Meechan, K. D. Micheva, K. Narayan, C. Pape, R. G. Parton, N. L. Schieber, Y. Schwab, B. Titze, P. Verkade, A. Weigel, L. M. Collinson, Nat Rev Methods Primers 2022, 2, 51.

D. P. Hoffman, G. Shtengel, C. S. Xu, K. R. Campbell, M. Freeman, L. Wang, D. E. Milkie, H. A. Pasolli, N. Iyer, J. A. Bogovic, D. R. Stabley, A. Shirinifard, S. Pang, D. Peale, K. Schaefer, W. Pomp, C.-L. Chang, J. Lippincott-Schwartz, T. Kirchhausen, D. J. Solecki, E. Betzig, H. F. Hess, Science 2020, 367, eaaz5357.

M. Januszewski, J. Kornfeld, P. H. Li, A. Pope, T. Blakely, L. Lindsey, J. Maitin-Shepard, M. Tyka, W. Denk, V. Jain, Nat Methods 2018, 15, 605.

X. Chen, C. Qiao, T. Jiang, J. Liu, Q. Meng, Y. Zeng, H. Chen, H. Qiao, D. Li, J. Wu, PhotoniX 2024, 5, 4.

A. Saguy, O. Alalouf, N. Opatovski, S. Jang, M. Heilemann, Y. Shechtman, Nat Methods 2023, 20, 1939.

C. Qiao, S. Liu, Y. Wang, W. Xu, X. Geng, T. Jiang, J. Zhang, Q. Meng, H. Qiao, D. Li, Q. Dai, Nat Biotechnol 2025.

E. D. Zhong, T. Bepler, B. Berger, J. H. Davis, Nat Methods 2021, 18, 176.

S. Aiyer, P. R. Baldwin, S. M. Tan, Z. Shan, J. Oh, A. Mehrani, M. E. Bowman, G. Louie, D. O. Passos, S. Đorđević-Marquardt, M. Mietzsch, J. A. Hull, S. Hoshika, B. A. Barad, D. A. Grotjahn, R. McKenna, M. Agbandje-McKenna, S. A. Benner, J. A. P. Noel, D. Wang, Y. Z. Tan, D. Lyumkis, Nat Commun 2024, 15, 389.

I. Lazić, M. Wirix, M. L. Leidl, F. De Haas, D. Mann, M. Beckers, E. V. Pechnikova, K. Müller-Caspary, R. Egoavil, E. G. T. Bosch, C. Sachse, Nat Methods 2022, 19, 1126.

M. Moor, O. Banerjee, Z. S. H. Abad, H. M. Krumholz, J. Leskovec, E. J. Topol, P. Rajpurkar, Nature 2023, 616, 259.

C. Ma, W. Tan, R. He, B. Yan, Nat Methods 2024, 21, 1558.

A. Kondepudi, M. Pekmezci, X. Hou, K. Scotford, C. Jiang, A. Rao, E. S. Harake, A. Chowdury, W. Al-Holou, L. Wang, A. Pandey, P. R. Lowenstein, M. G. Castro, L. I. Koerner, T. Roetzer-Pejrimovsky, G. Widhalm, S. Camelo-Piragua, M. Movahed-Ezazi, D. A. Orringer, H. Lee, C. Freudiger, M. Berger, S. Hervey-Jumper, T. Hollon, Nature 2025, 637, 439.

R. J. Chen, T. Ding, M. Y. Lu, D. F. K. Williamson, G. Jaume, A. H. Song, B. Chen, A. Zhang, D. Shao, M. Shaban, M. Williams, L. Oldenburg, L. L. Weishaupt, J. J. Wang, A. Vaidya, L. P. Le, G. Gerber, S. Sahai, W. Williams, F. Mahmood, Nat Med 2024, 30, 850.

R. Rombach, A. Blattmann, D. Lorenz, P. Esser, B. Ommer, in Proceedings of the IEEE/CVF Conference on Computer Vision and Pattern Recognition (CVPR), 2022, pp. 10684–10695.

B. Zhou, A. Khosla, A. Lapedriza, A. Oliva, A. Torralba, in 2016 IEEE Conference on Computer Vision and Pattern Recognition (CVPR), IEEE, Las Vegas, NV, USA 2016, pp. 2921–2929.

J. J. P. Peters, T. Mullarkey, E. Hedley, K. H. Müller, A. Porter, A. Mostaed, L. Jones, Nat Commun 2023, 14, 5184.

C. Lu, K. Chen, H. Qiu, X. Chen, G. Chen, X. Qi, H. Jiang, Nat Commun 2024, 15, 4677.

U. Ermel, A. Cheng, J. X. Ni, J. Gadling, M. Venkatakrishnan, K. Evans, J. Asuncion, A. Sweet, J. Pourroy, Z. S. Wang, K. Khandwala, B. Nelson, D. McCarthy, E. M. Wang, R. Agarwal, B. Carragher, Nat Methods 2024, 21, 2200.

K. J. Czymmek, I. Belevich, J. Bischof, A. Mathur, L. Collinson, E. Jokitalo, Nat Cell Biol 2024, 26, 498.

C. Stringer, M. Pachitariu, Nat Methods 2025, 22, 592.

X. Feng, Nature Plants 2023, 9.

K. M. Harris, J. Spacek, M. E. Bell, P. H. Parker, L. F. Lindsey, A. D. Baden, J. T. Vogelstein, R. Burns, Sci Data 2015, 2, 150046.

N. Kasthuri, K. J. Hayworth, D. R. Berger, R. L. Schalek, J. A. Conchello, S. Knowles-Barley, D. Lee, A. Vázquez-Reina, V. Kaynig, T. R. Jones, M. Roberts, J. L. Morgan, J. C. Tapia, H. S. Seung, W. G. Roncal, J. T. Vogelstein, R. Burns, D. L. Sussman, C. E. Priebe, H. Pfister, J. W. Lichtman, Cell 2015, 162, 648.

M. R. Wozny, A. Di Luca, D. R. Morado, A. Picco, R. Khaddaj, P. Campomanes, L. Ivanović, P. C. Hoffmann, E. A. Miller, S. Vanni, W. Kukulski, Nature 2023, 618, 188.

W. Zhao, S. Zhao, Z. Han, X. Ding, G. Hu, L. Qu, Y. Huang, X. Wang, H. Mao, Y. Jiu, Y. Hu, J. Tan, X. Ding, L. Chen, C. Guo, H. Li, Nat. Photon. 2023, 17, 806.

J. Engelmann, M. O. Bernabeu, Nat Commun 2025, 16, 6862.

J. R. Kremer, D. N. Mastronarde, J. R. McIntosh, Journal of Structural Biology 1996, 116, 71.

L. Xie, X. Song, Z. Liao, B. Wu, J. Yang, H. Zhang, J. Hong, Micron 2019, 120, 80.

K. Zheng, C. Lu, J. Chen, J. Zhu, Advances in Neural Information Processing Systems 2023, 36, 55502.

B. Kawar, M. Elad, S. Ermon, J. Song, in Advances in Neural Information Processing Systems (Eds.: S. Koyejo, S. Mohamed, A. Agarwal, D. Belgrave, K. Cho, A. Oh), Vol. 35, Curran Associates, Inc. 2022, pp. 23593–23606.

S. W. Zamir, A. Arora, S. Khan, M. Hayat, F. S. Khan, M.-H. Yang, in 2022 IEEE/CVF Conference on Computer Vision and Pattern Recognition (CVPR), IEEE, New Orleans, LA, USA 2022, pp. 5718–5729.

