## Supplementary Information for "A foundation AI model enhances electron microscopy image analysis"

This file contains Supplementary Methods, Supplementary Figures, Supplementary Tables and Supplementary References.

### Supplementary Figures

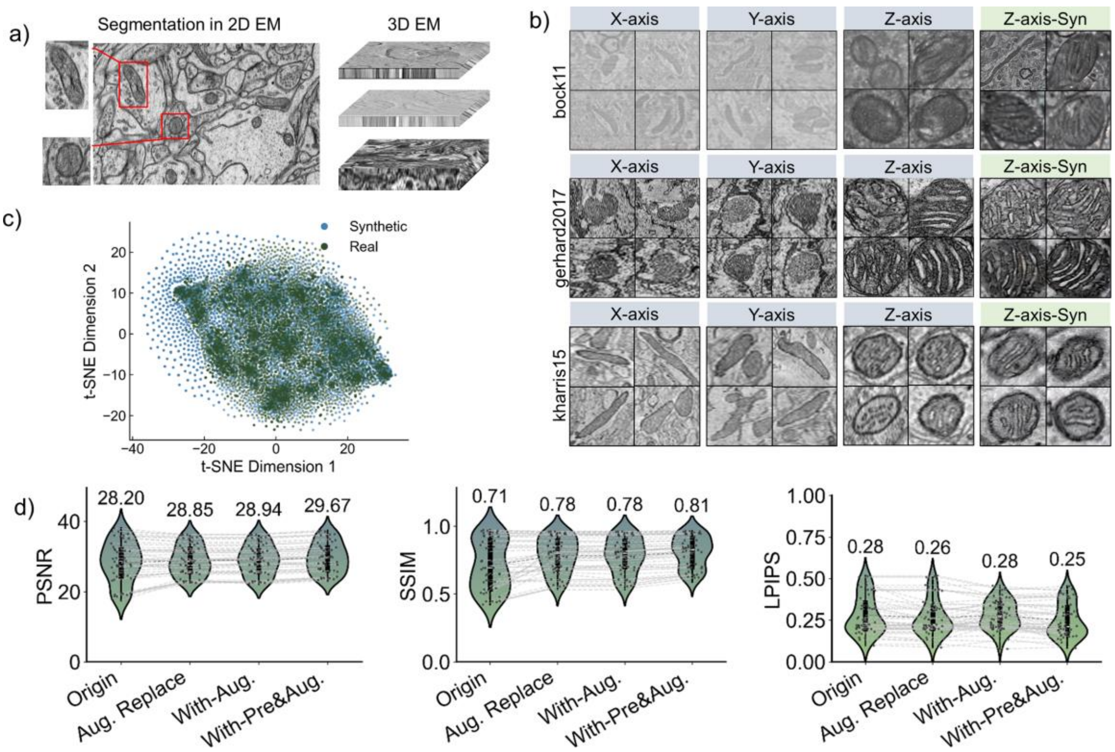


**Supplementary Fig. 1 Construction of a multi-source dataset - MemEM.** a) High- and low-resolution membrane images are obtained from real two-dimensional (2D) and three-dimensional (3D) electron microscopy (EM) data, where 3D EM includes volume electron microscopy (vEM) and electron tomography (ET). b) Examples of enhanced EM membrane images. Columns 1 and 2 show real membrane segmented from the X- and Y-planes, column 3 displays real z-plane segments, and column 4 presents enhanced z-plane counterparts. c) t-SNE clustering analysis of real vs. enhanced data. Features obtained via Histogram of Oriented Gradients (HOG) are normalized and then clustered using t-SNE for dimensionality reduction analysis. We analyzed 6,000 real and 6,000 enhanced images (n = 12,000). d) Performance evaluation of enhanced data and our preprocessing method in super-resolution tasks. “Aug Replace” denotes replacing 500K real images with enhanced ones, yielding a mixed training set (1M real + 500K enhanced). Benchmarking was performed on the super-resolution validation set (Extended Data Table 2).

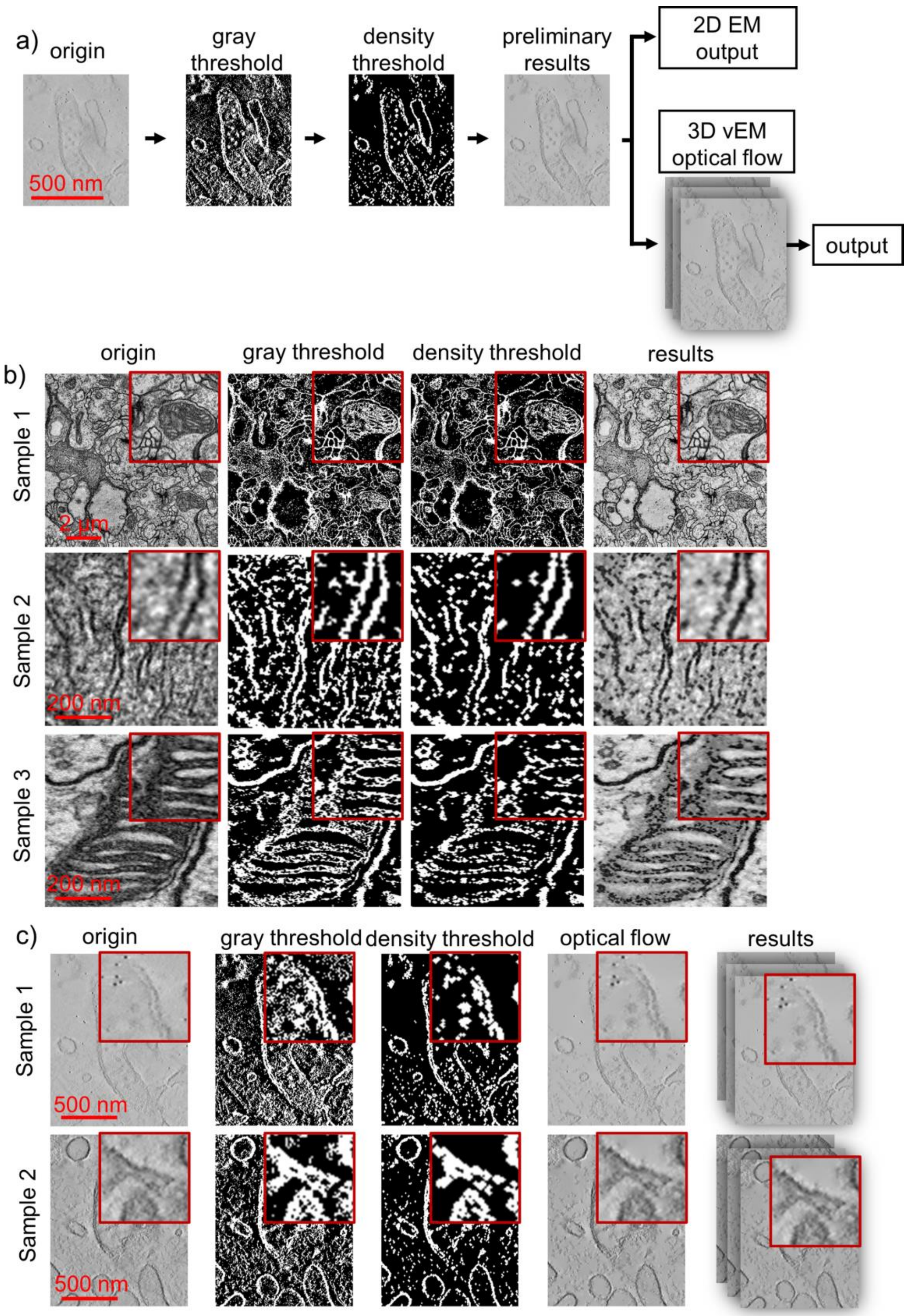


**Supplementary Fig. 2 Image preprocessing.** a) High-resolution membrane images from animal cell lines (e.g., mouse myoblasts) are used as input. Grayscale thresholding isolates dark regions (top 30% intensity pixels), followed by local density thresholding (top 70% density in a 3×3 window) to outline coarse membrane structures. For 2D images, results are directly output. For 3D microscopy data, optical flow estimation using the Farneback algorithm tracks pixel motion across adjacent slices to reconstruct missing membrane details, producing continuous 3D membrane contours. b) 2D image preprocessing. Large-scale or low-resolution membrane images undergo grayscale thresholding to extract membrane regions, followed by density thresholding to enhance boundary clarity. Contrast-limited adaptive histogram equalization and non-local means denoising sharpen membrane edges and improve image quality. c) 3D EM preprocessing. ET from mouse myoblast cell lines undergo 3D preprocessing. On the basis of 2D preprocessing, the missing 3D membrane details are reconstructed through optical flow estimation to ensure structural continuity.

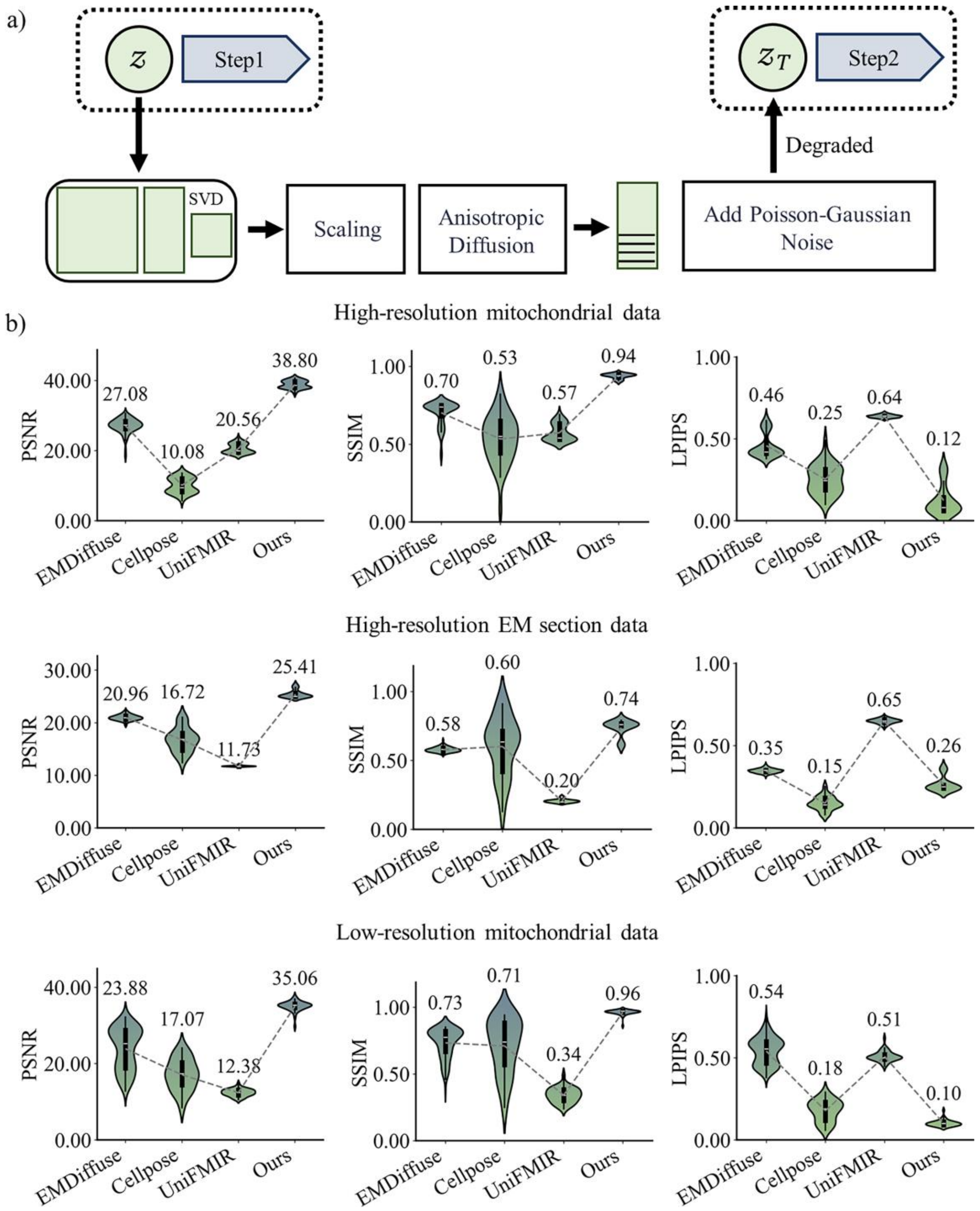


**Supplementary Fig. 3 Implementation of the denoising task and image quality assessment results.** a) Degradation method for the denoising task. Singular value decomposition (SVD) models diffusion and frequency attenuation, adding Poisson and Gaussian noise, with pseudoinverse operations to reverse degradation and recover clear images. b) Comparison of image quality metrics reconstructed by EMDiffuse1, Cellpose2, and UNiFMIR3 and DF5T. Denoising task validation sets were constructed from three data types (Supplementary Table 2).

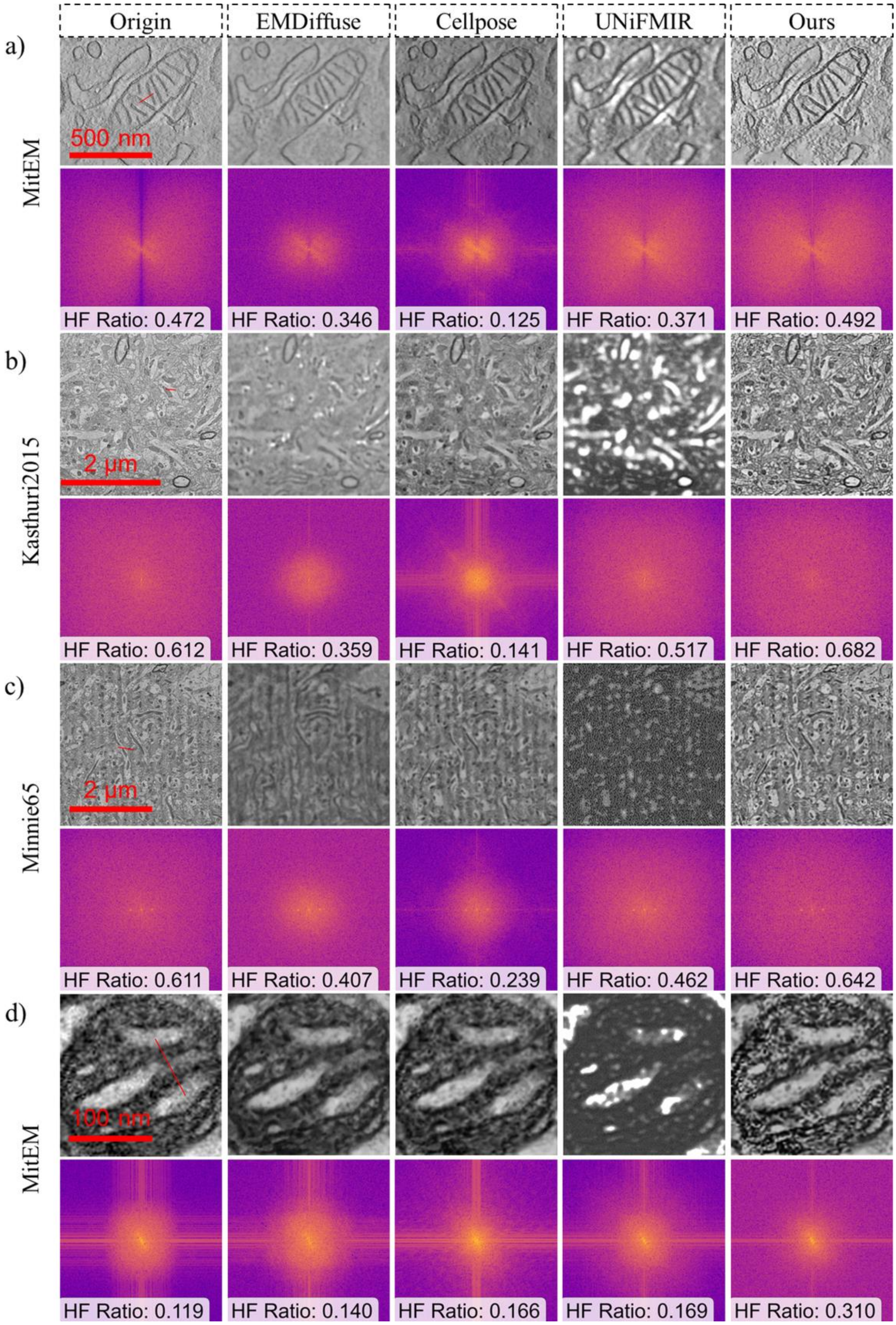


**Supplementary Fig. 4 Visualization of denoising task results and Fourier spectrum analysis.** Higher HF values indicate superior retention of high-frequency details. Among the four datasets, EMDiffuse and Cellpose exhibit weaker denoising performance, as evidenced by lower HF metrics, while UNiFMIR outputs show significant distortion and substantial loss of high-frequency details. In contrast, our method achieves optimal performance across all metrics, demonstrating exceptional detail recovery, superior denoising efficacy, and maximal preservation of high-frequency information.

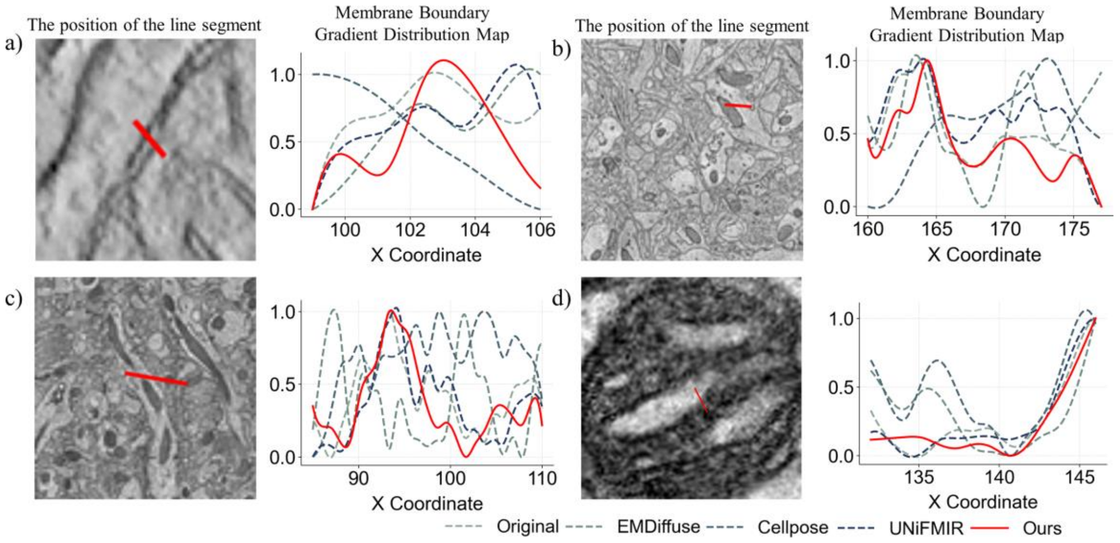


**Supplementary Fig. 5 Membrane boundary gradient distribution results for the denoising task from EMDiffuse, Cellpose, UNiFMIR, and DF5T.** The X Coordinate denotes the pixel position along the membrane boundary. Our method achieves the optimal balance between denoising efficacy and peak position alignment, demonstrating DF5T's cross-sample stability in managing noise suppression and structural preservation.

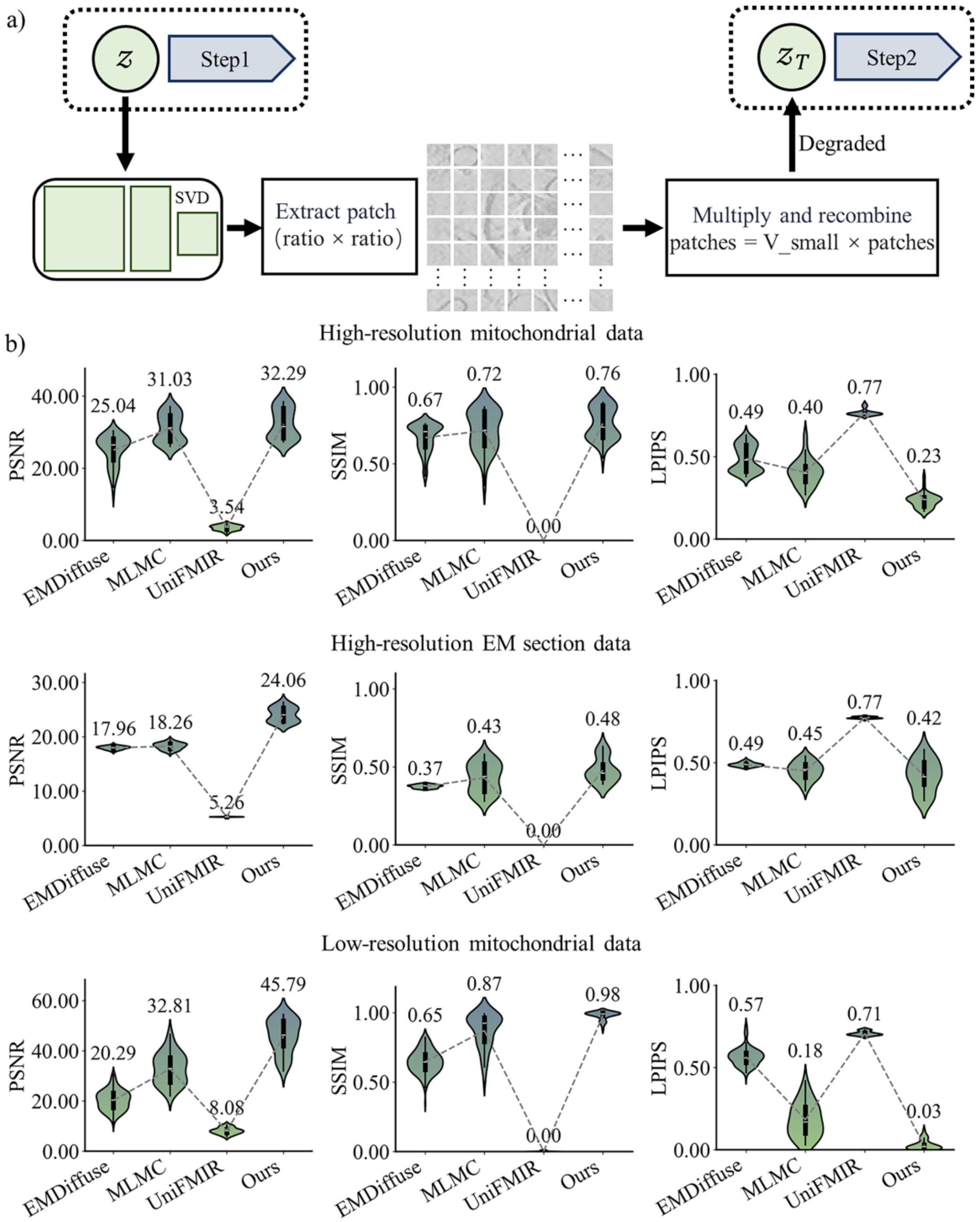


**Supplementary Fig. 6 Implementation of the super-resolution task and image quality assessment results.** a) Degradation method for the super-resolution task. SVD decomposes the downsampling matrix, combining pseudoinverse and bilinear interpolation to upscale low-resolution images, restoring high-resolution textures and edges. b) Comparison of image quality metrics reconstructed by EMDiffuse[1], MLMC[2], UNiFMIR[3] and DF5T. Super-resolution task validation sets were derived from the same generalization datasets (Supplementary Table 2).

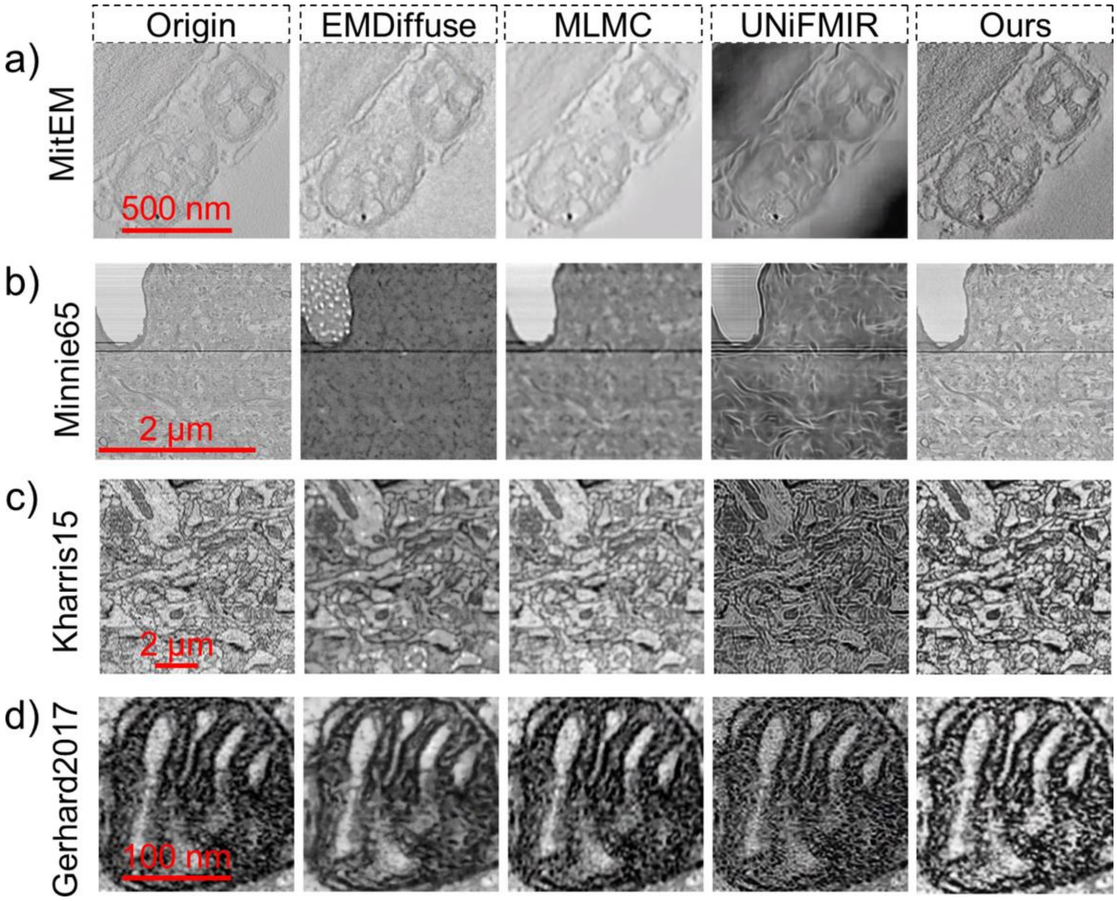


**Supplementary Fig. 7 Visualization of super-resolution task results reconstructed from EMDiffuse, MLMC, UNiFMIR, and DF5T across four datasets.** Our model delivers the best super-resolution effect, with sharper image details and texture/structure recovery closer to the origin.

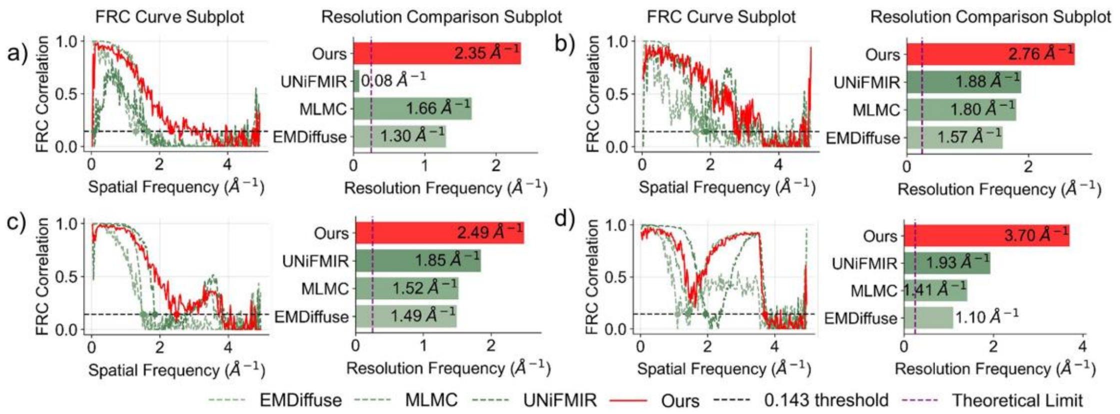


**Supplementary Fig. 8 Fourier ring correlation analysis (FRC) for the super-resolution task.** This figure presents FRC analysis results for the four models corresponding to Supplementary Fig. 7. Our method achieves the highest resolution frequency across all datasets, significantly outperforming others, and maintains a consistently high FRC curve level.

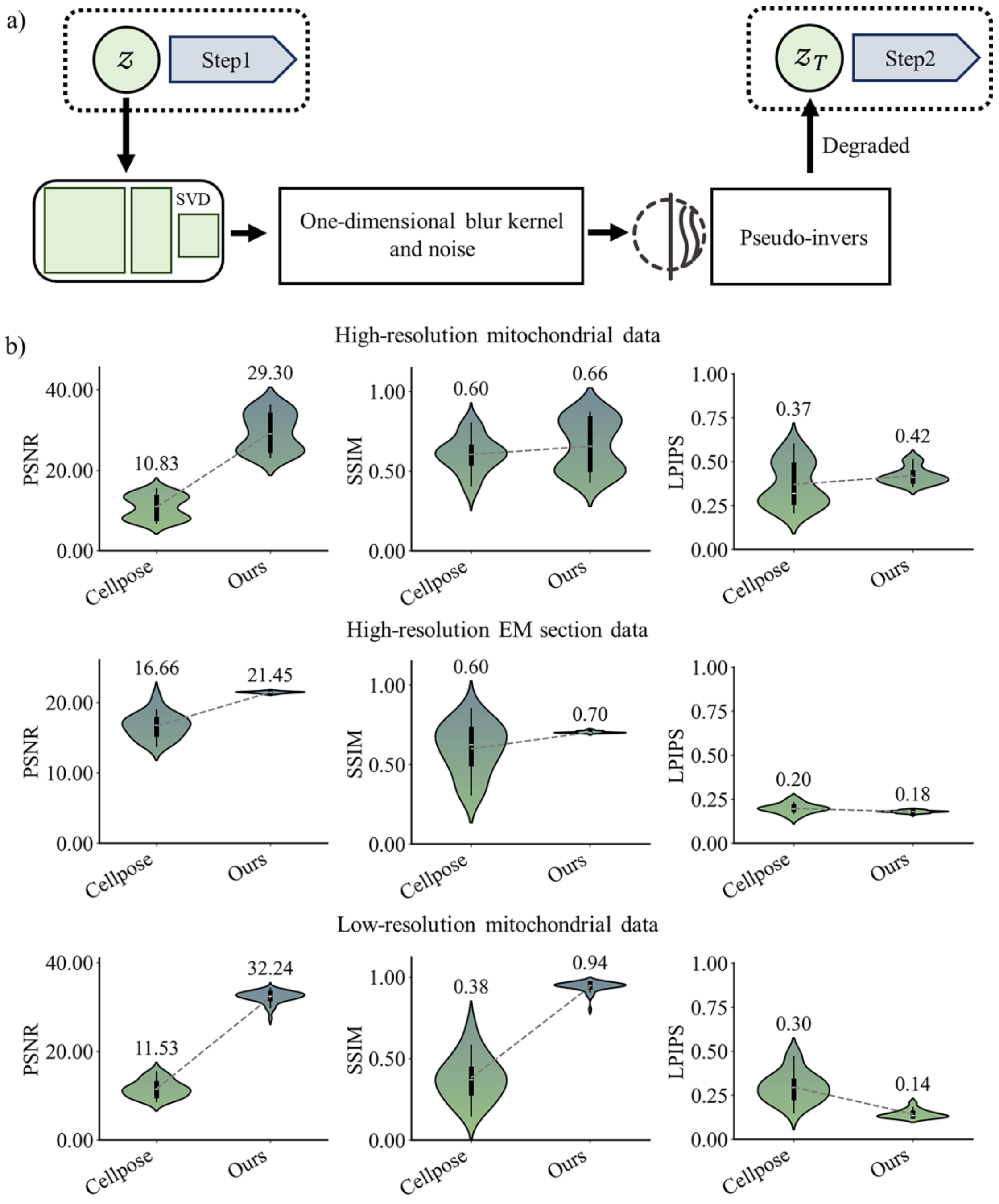


**Supplementary Fig. 9 Implementation of the deblurring task and image quality assessment results.** a) Degradation method for the deblurring task. SVD decomposes the 1D blur kernel matrix, applying regularized pseudoinverse to invert convolution blur, reconstructing high-frequency image details. b) Comparison of image quality metrics reconstructed by Cellpose[4] and DF5T. Deblurring task validation sets were built from the same generalization datasets (Supplementary Table 2).

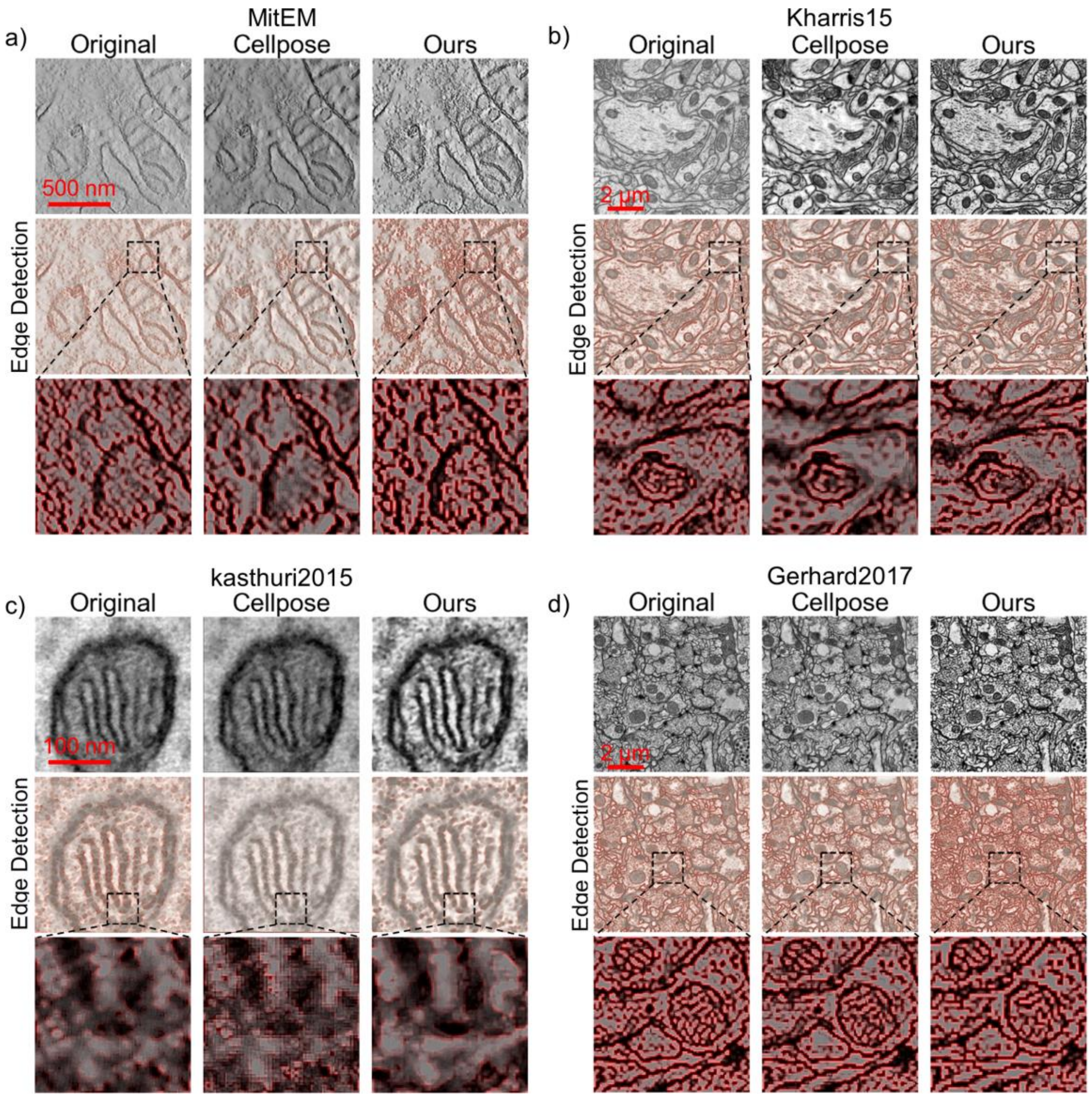


**Supplementary Fig. 10 Visualization of the deblurring task.** This figure compares the performance of Cellpose and DF5T in edge sharpness enhancement across four representative images. For each sample, the first row shows the raw image, the second row displays edge detection results, and the third row presents local enhancement results. Our method's red gradient edges in local enhancement exhibit greater sharpness and contrast, indicating superior deblurring performance.

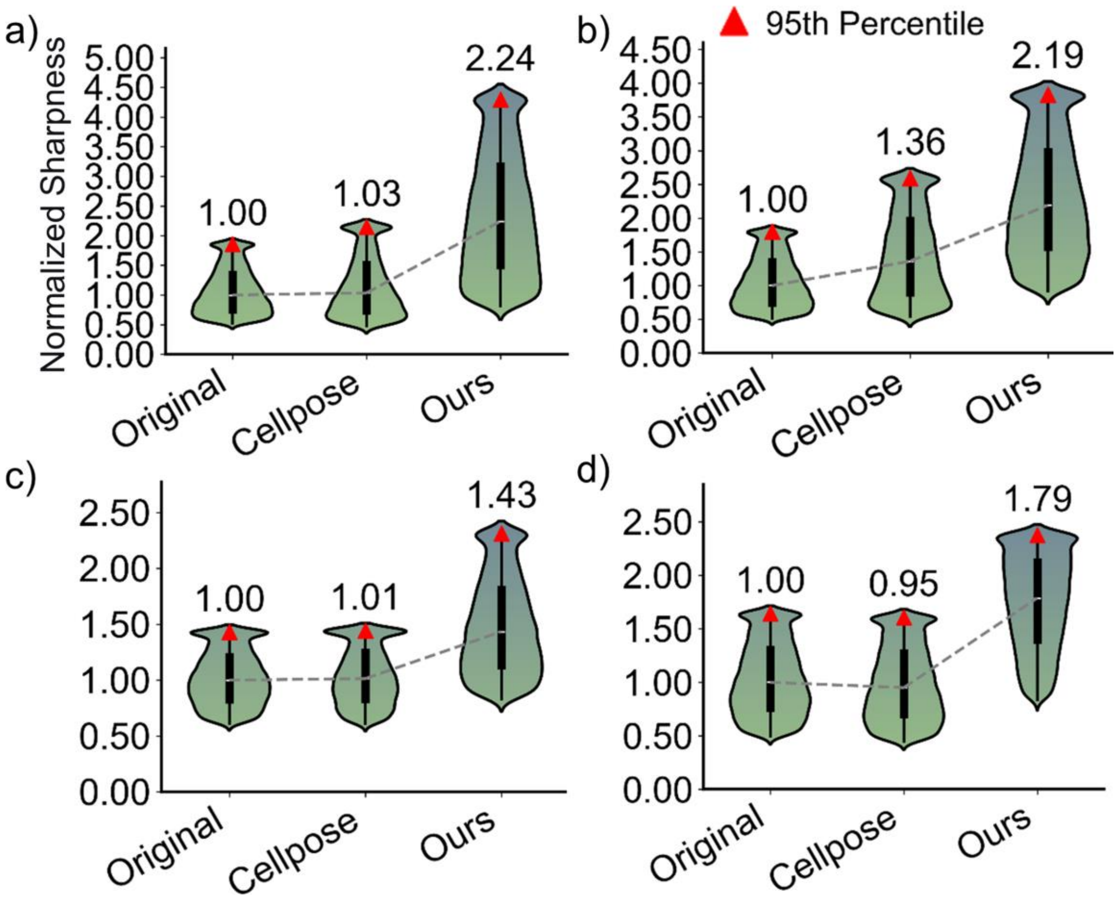


**Supplementary Fig. 11 Comparison of normalized edge sharpness for deblurring results.** This figure presents normalized edge sharpness comparisons corresponding to Supplementary Fig. 10. Our method outperforms Cellpose and the original images in median sharpness and 95th percentile, with a more concentrated and higher sharpness distribution, achieving optimal clarity and stability in edge enhancement, particularly in complex membrane regions.

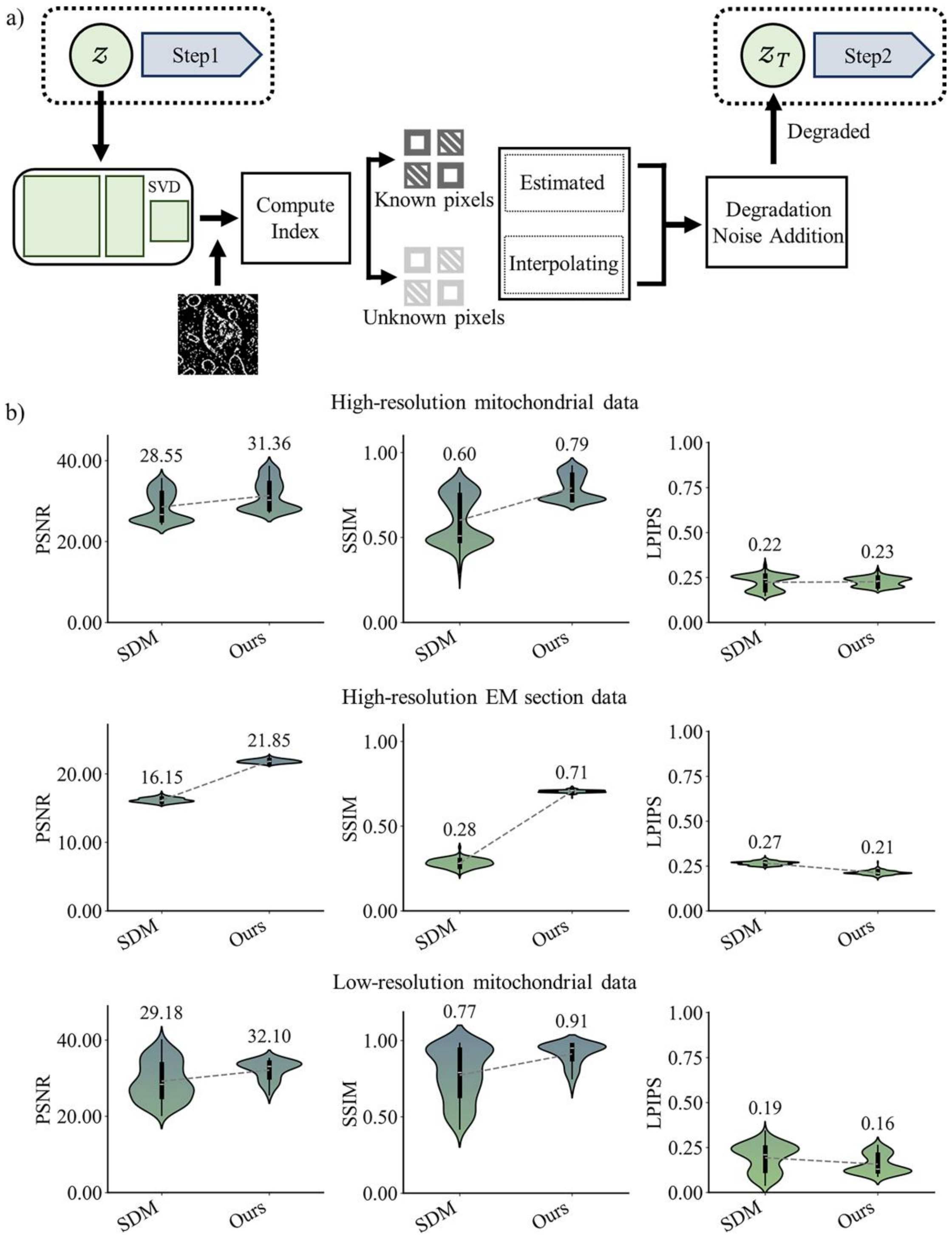


**Supplementary Fig. 12 Implementation of the 2D inpainting task and image quality assessment results.** a) Degradation method for the 2D inpainting task. SVD decomposes the mask matrix to exclude missing pixels, using 2D Gaussian kernel smoothing to accurately restore structure and texture in image gaps. b) Comparison of image quality metrics reconstructed by SDM and DF5T. 2D inpainting task validation sets were constructed from the same generalization datasets (Supplementary Table 2).

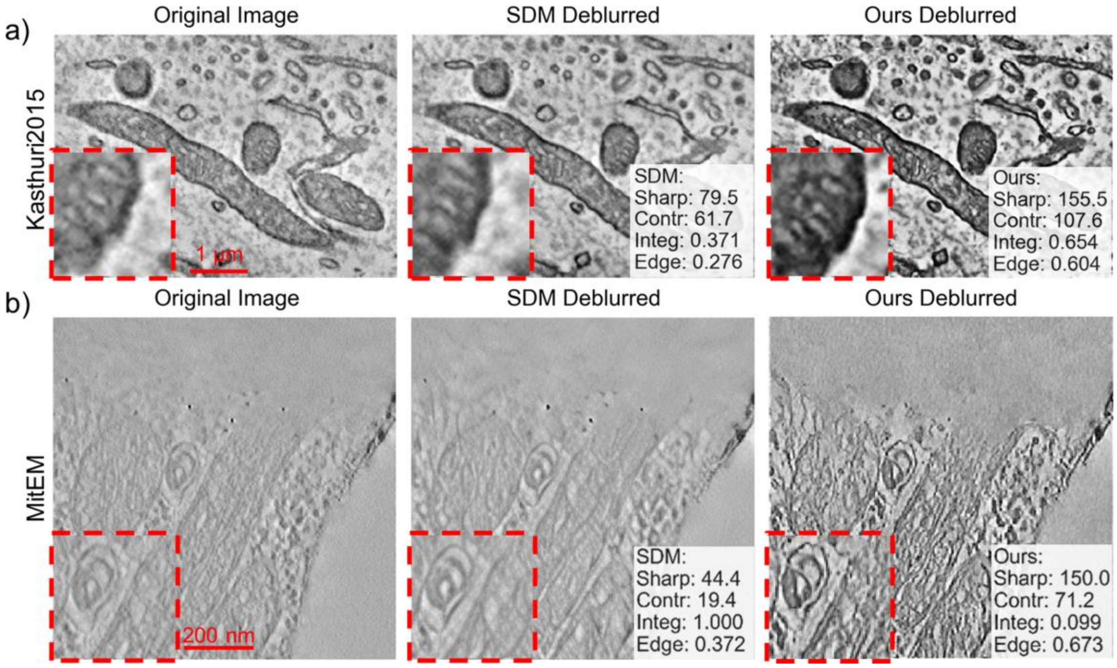


**Supplementary Fig. 13 Representative 2D inpainting images reconstructed by SDM and DF5T.** Performance is quantitatively assessed by membrane sharpness, organelle contrast, structural integrity, and edge preservation, where superior scores denote enhanced image quality.

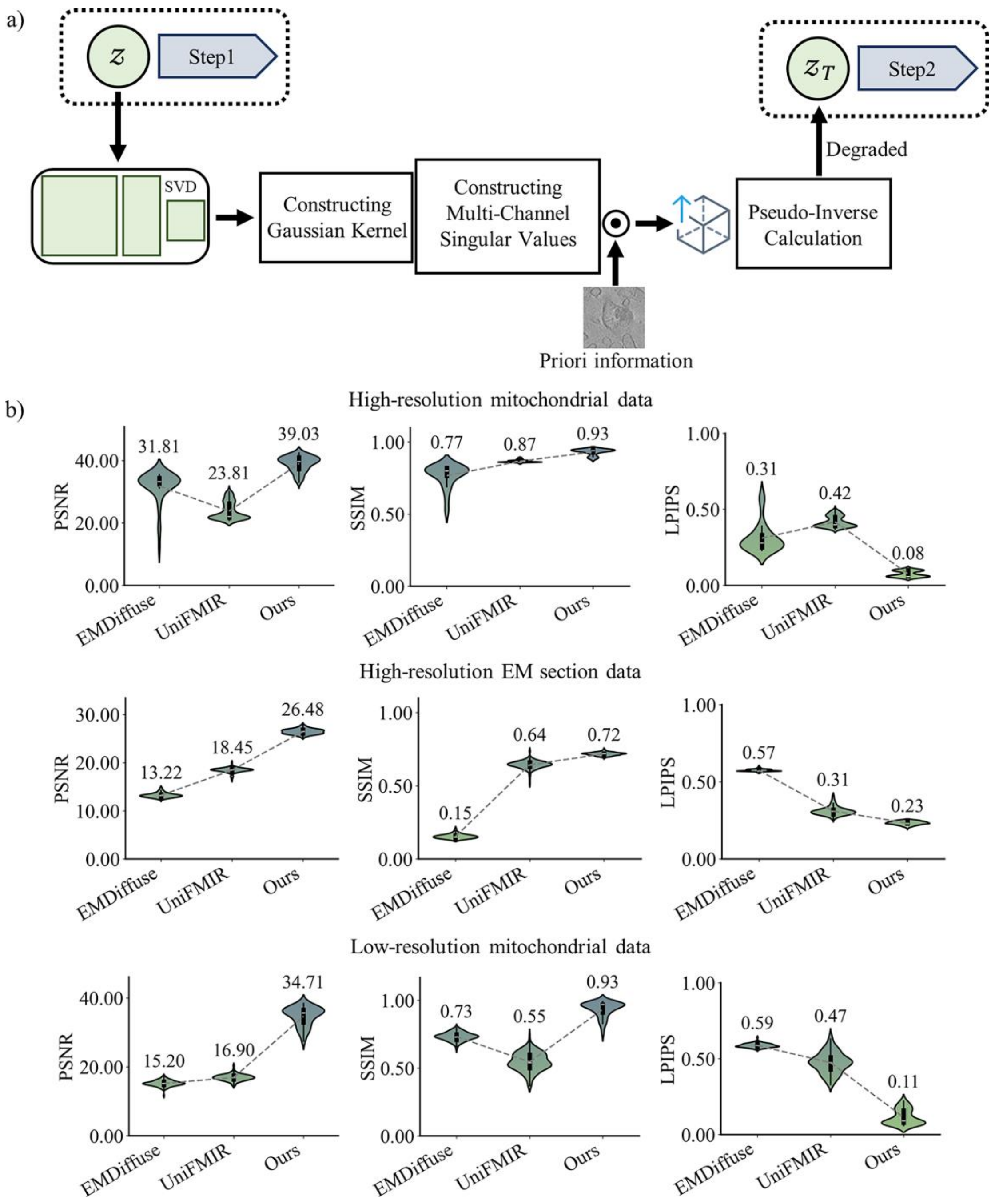


**Supplementary Fig. 14 Implementation of the 3D isotropic restoration task and image quality metrics.** a) SVD decomposes a 3D Gaussian kernel, integrating prior frame features with SSIM weights to enhance temporal consistency and optimize image restoration details. b) Comparison of image quality metrics reconstructed by EMDiffuse, UNiFMIR and DF5T. 3D isotropic restoration validation sets were built from the same generalization datasets (Supplementary Table 2).

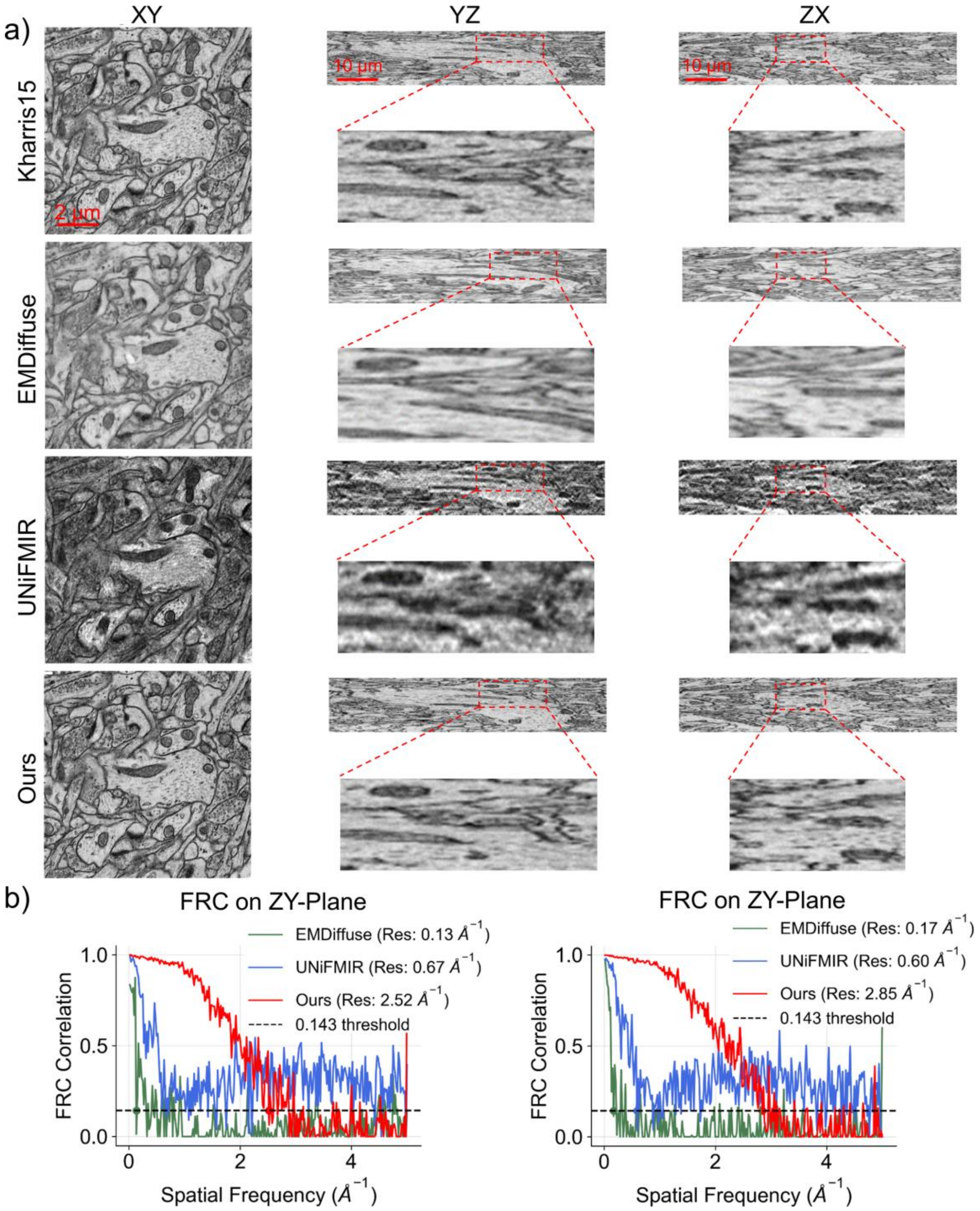


**Supplementary Fig. 15 Visualization of 3D isotropic restoration results for the provided dataset.** a) Representative orthogonal central slice images (XY, YZ, ZX) reconstructed from EMDiffuse, UNiFMIR and DF5T. b) FRC curves on the ZY and ZX planes show that our method's correlation decays more slowly, achieving higher resolution limits than EMDiffuse and UNiFMIR, indicating superior preservation of high-frequency details.

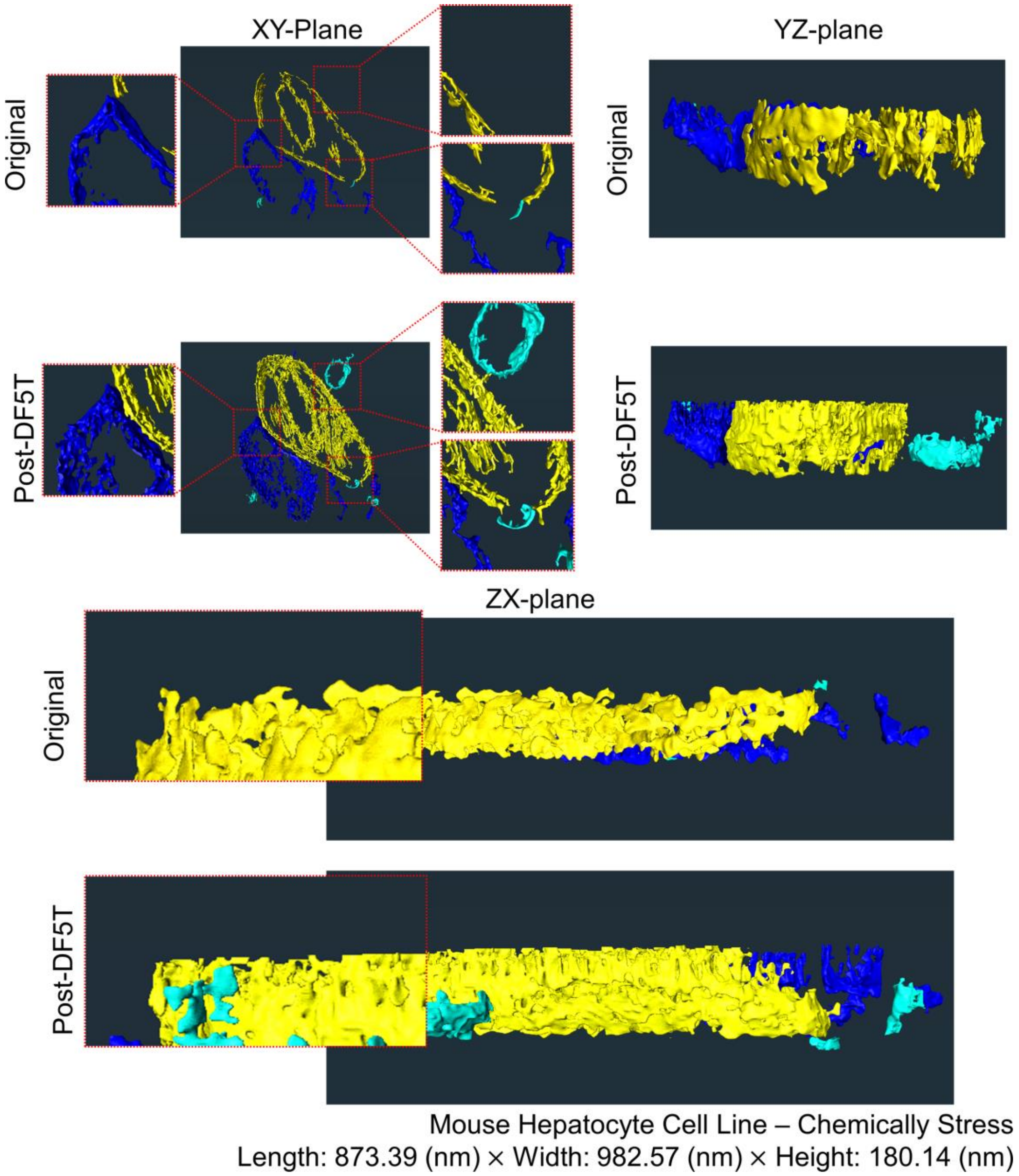


**Supplementary Fig. 16 3D reconstruction of chemically stressed mouse hepatocyte cell line mitochondria before and after DF5T treatment.** Following treatment, the adhesion between neighboring mitochondria is significantly reduced, allowing for complete segmentation of individual mitochondria. Additionally, the mitochondria and their internal membrane structures appear more intact and smoother.

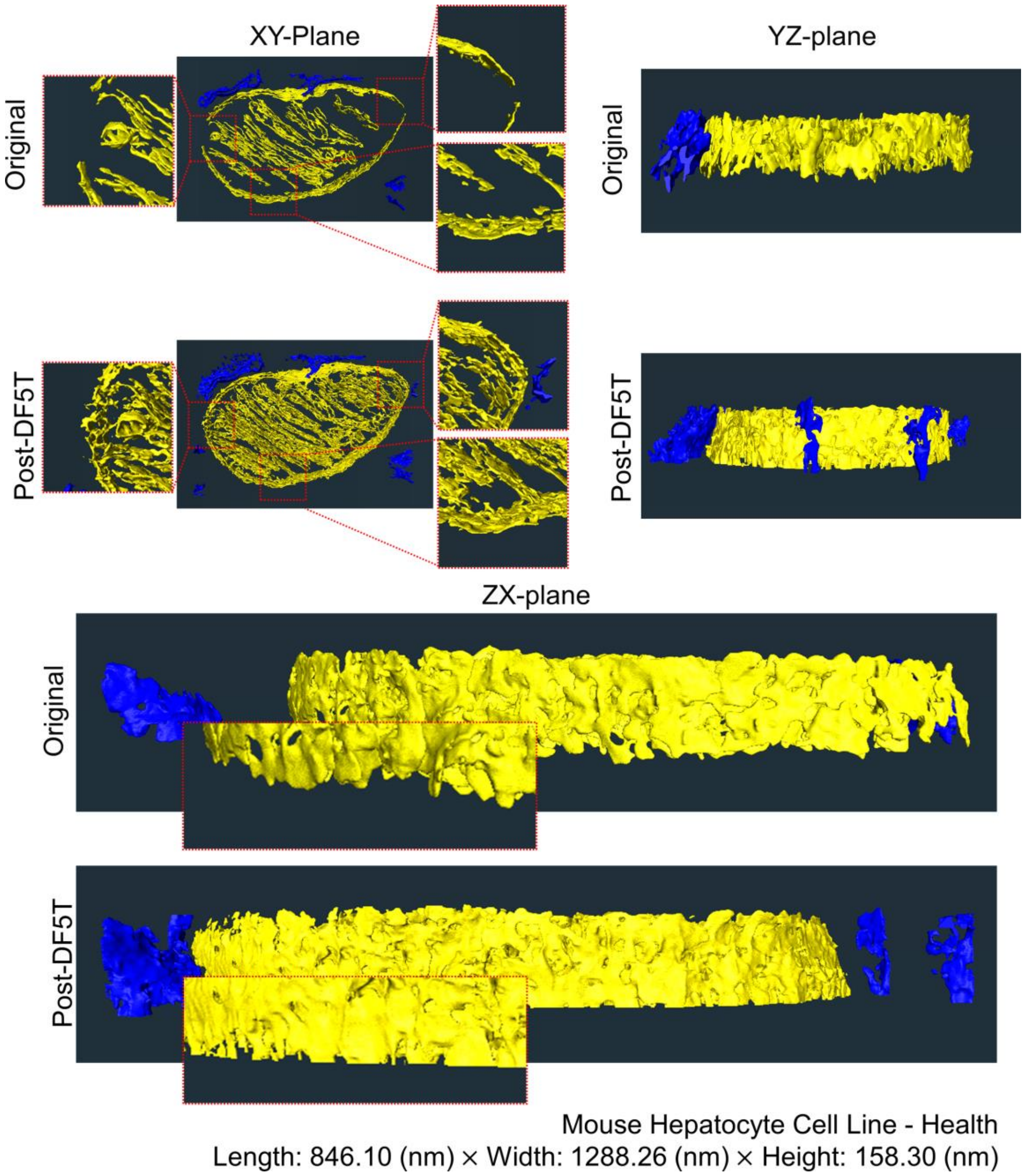


**Supplementary Fig. 17 3D reconstruction of healthy mouse hepatocyte cell line mitochondria before and after DF5T treatment.** Corresponding to Supplementary Fig. 16’s slice, healthy results show increased mitochondrial cristae region. enhancing segmentation, and damaged membrane structures from the “Before” state are restored to complete membranes.

### Supplementary Table

**Supplementary Table 1 Information on datasets involved in training.** This table details sources of EM data and public image datasets.

| Dataset name | Introduction | Country | Amount of data | Cropped images(slices) | Number of Membrane-bound organelles segmented(Nucleus, mitochondria, Golgi, chloroplasts, lysosomes) | Source | Dataset Size | Imaging Techniques |
|---|---|---|---|---|---|---|---|---|
| CEM1.5M | mitochondria | America | | 1500000 | | https://www.ebi.ac.uk/empiar/EMPIAR-11035/ | | |
| CEM500k | large-scale heterogeneous unlabeled cellular electron microscopy image | America | | | | https://www.ebi.ac.uk/empiar/EMPIAR-10592/ | | |
| The Plantorgan hunter dataset | 19 kinds of plants | China | | 3233 | 3233 mitochondria and Chloroplast | https://cstr.cn/31253.11.sciencedb.01335 | z 2μm | Transmission Electron Microscopy |
| Phelps_hildebrand_graham2021 | the ventral nerve cord of an adult female Drosophila melanogaster | America | data = channel[1228:1229, 115965:116989, 23124:24148] | 115956 | 165591 mitochondria | https://bossdb.org/project/phelps_hildebrand_graham2021 | x/y 4.3nm, z 45nm | FEI Spirit TEM |
| Minnie65_8x8x40 | cortical circuits within the visual cortex of mouse | America | data = channel[19000:19016, 56298:57322, | 82890 | 192465 mitochondria and | https://bossdb.org/project/microns-minnie | x/y 8nm, z 40nm | Serial Block-Face Scanning Electron Microscopy |
| Gerhard2017 | 1st and 3rd instar Drosophila melanogaster larvae | America | data = channel[206:210, 19843:20843, 27889:28889] | 120005 | 242233 mitochondria | https://doi.org/10.60533/BOSS-2017-0DCV | x/y 2.3nm, z 50nm | FEI Spirit TEM |
| Bock11 | mouse primary visual cortical data | America | data = channel[3000:3003, 60000:62048, 67634:69690] | 138240 | 289224 mitochondria | https://doi.org/10.60533/BOSS-2011-K12H | x/y 4nm, z 40nm | X-ray crystallography |
| Kharris15 | hippocampal neuropil | America | data = channel[100:110, 4096:4608, 4096:4608] | 6240 | 1743 mitochondria | https://doi.org/10.60533/BOSS-2015-GYEN | x/y 2nm, z 50nm | Focused Ion Beam Scanning Electron Microscopy |
| Kasthuri2015 | mouse cortex | America | data = channel[900:910, 7000:8000, 5000:6000] | 15526 | 51771 mitochondria | https://doi.org/10.60533/BOSS-2015-IOQM | x/y 6nm, z 30nm | Serial Block-Face Scanning Electron Microscopy |
| EMPIAR-12482 | Imod prealign files of synaptosomes isolated without protease inhibitor or DTT | America | data = channel[0:40, 0:5760, 0:4092] | 3596 | 3596 synaptosomes | https://www.ebi.ac.uk/empiar/EMPIAR-12482/ | (3.3 Å, 3.3 Å) | Cryo-electron tomography |
| | Imod prealign files of synaptosomes isolated with protease inhibitor and DTT | America | data = channel[0:31, 0:5760, 0:4092] | 2787 | 2787 synaptosomes | | | |
| jrc_mus-liver-7 | Obese Male Mouse Liver | America | data = channel[0:3584, 0:8192, 0:8192] | 102400 | | https://openorganelle.janelia.org/datasets/jrc_mus-liver-7 | x/y 8nm, z 8nm | Focused Ion Beam Scanning Electron Microscopy |
| EMPIAR-11589 | Cryo-Electron Tomography of Golgi Apparatus from a Zebra Fish Embryo | America | data = channel[0:1, 0:3710, 0:3838] | 54 | 54 Golgi | https://www.ebi.ac.uk/empiar/EMPIAR-11589/ | UNSIGNED 16 BIT INTEGER | Cryo-Electron Tomography |
| NucMM | Zebrafish brain | America | data = channel[0:397, 0:1450, 0:2000] | 397 | 132896 Nucleus | https://opendatalab.org.cn/OpenDataLab/NucMM | x/y 0.51μm, z 0.48μm | Serial Block-Face Scanning Electron Microscopy |
| | Mouse visual cortex | | data = channel[0:700, 0:996, 0:968] | 192 | 20322 Nucleus | | | |
| JRCData-allone-1024 | ic_desmosome (Intercellular desmosome) | America | | 16920 | | https://open.quiltdata.com/b/janelia-cosem-datasets/tree/ | x/y 4nm, z 45nm | Focused Ion Beam Scanning Electron Microscopy |
| | jr_cos (COS-7 cell) | | | 3668 | | | x/y 4nm, z 50nm | |
| | jrc_ctl-id8 (ctl-id8 cell line) | | | 58159 | | | x/y 4nm, z 40nm | |
| | jr_dauer-larva(C. elegans dauer larva (whole organism)) | | | 7858 | | | x/y 4nm, z 50nm | |
| | jr_fly-fsb-1 (Drosophila flight muscle (FSB cell)) | | | 57565 | | | x/y 4nm, z 45nm | |
| | jr_hela (HeLa cell (human cervical cancer)) | | | 49945 | | | x/y 4nm, z 40nm | |
| | jr_hela-h89 (HeLa cell) | | | 11519 | | | x/y 4nm, z 45nm | |
| | jr_jurkat （Jurkat cell (human T-lymphocyte)） | | | 7297 | | | x/y 4nm, z 40nm | |
| | jr_mus-epididymis （Mouse epididymis tissue） | | | 11649 | | | x/y 4nm, z 50nm | |
| | jr_mus-guard-hair-follicle （Mouse guard hair follicle） | | | 15037 | | | x/y 4nm, z 45nm | |
| | jr_mus-kidney （Mouse kidney tissue） | | | 44760 | | | x/y 4nm, z 50nm | |
| EMPIAR-11746 | FIB-SEM dataset of a human bone osteosarcoma epithelial cell (U2-OS) | Finland | data = channel[0:1168, 0:3394, 0:1385] | 7426 | 1097 Nuclear envelope,781 Lipid droplets,516 Lysosomes,1168 Golgi | https://www.ebi.ac.uk/empiar/EMPIAR-11746/ | x/y 2.5nm, z 5nm | Focused Ion Beam Scanning Electron Microscopy |
| EMPIAR-11849 | FIB-SEM dataset showing localization of a Golgi matrix protein GM130 in human hepatocellular carcinoma cell (Huh-7) | Finland | data = channel[0:564, 0:3269, 0:1397] | 564 | 564 Golgi | https://www.ebi.ac.uk/empiar/EMPIAR-11849/ | (40 Å, 40 Å) | Focused Ion Beam Scanning Electron Microscopy |
| EMPIAR-11819 | Cryo-electron tomogram of mitochondria in cryo-FIB milled l-Opa1* mouse embryonic fibroblasts | America | data = channel[0:245, 0:11520, 0:8184] | 1440 | 1440 mitochondria | https://www.ebi.ac.uk/empiar/EMPIAR-11819/ | (1.375 Å, 1.375 Å) | Cryo-Electron Tomography |
| Ours (real) | Healthy Mouse Myoblast Cell Line | China | | 998 | 998 | https://www.scidb.cn/detail?dataSetId=8d30c6b23acd46d09e44114e8f739fe4#p3 | z 0.54nm | Transmission Electron Microscopy |
| | Diseased Mouse Myoblast Cell Line | | | 321 | 321 | | | |
| | Healthy Tobacco Mesophyll Cells | | | 251 | 251 | | | |
| | Tobacco Mesophyll Cells(Pore Formation in the Membrane via Molecular Biology Techniques) | | | 720 | 720 | | | |
| Ours (Synthesis) | Synthetic data (generated by SDM) | | | | 1000 Cryo-ET golgi, 10000 FIB-SEM golgi, 10000 Cryo-ET synaptosomes, 10000 TEM Chloroplast | | | |
| Total amount | | | | Total: 2.25 million images (organelles | | | | |

**Supplementary Table 2 Test datasets of DF5T.** This table outlines the composition of test sets for each task and describes testing methods.

| Task | Type | Count | Source of Data | Image quality evaluation metric | Task evaluation metric |
|---|---|---|---|---|---|
| Denoise | High-resolution organelle | 40 | Healthy Mouse Myoblast Cell Line | PSNR、SSIM and LPIPS | Evaluated using smoothed gradient profiles and Fourier spectrum analysis. |
| | Large-scale slice | 40 | Random non-repeated slices from six public electron tomography datasets. | | |
| | Low-resolution organelle | 40 | Randomly selected non-repeated segmentation results from slices of six public electron tomography datasets. | | |
| Super resolution | High-resolution organelle | 40 | Healthy Mouse Myoblast Cell Line | | Evaluated by FRC curves and resolution limits at a 0.143 threshold (Å$^{-1}$), compared to the theoretical diffraction limit. |
| | Large-scale slice | 40 | Random non-repeated slices from six public electron tomography datasets. | | |
| | Low-resolution organelle | 40 | Randomly selected non-repeated segmentation results from slices of six public electron tomography datasets. | | |
| Deblur | High-resolution organelle | 40 | Healthy Mouse Myoblast Cell Line | | Evaluated using edge sharpness metrics derived from trimmed gradient magnitudes at detected edges. |
| | Large-scale slice | 40 | Random non-repeated slices from six public electron tomography datasets. | | |
| | Low-resolution organelle | 40 | Randomly selected non-repeated segmentation results from slices of six public electron tomography datasets. | | |
| 2D Inpaint | High-resolution organelle | 40 | Healthy Mouse Myoblast Cell Line | | Evaluated using topological metrics (endpoint and branch point ratios), SSIM, and edge similarities (Canny and Sobel) |
| | Large-scale slice | 40 | Random non-repeated slices from six public electron tomography datasets. | | |
| | Low-resolution organelle | 40 | Randomly selected non-repeated segmentation results from slices of six public electron tomography datasets. | | |
| 3D Isotropic | High-resolution organelle | 40 | Healthy Mouse Myoblast Cell Line | | Evaluated by analyzing FFT magnitude profiles along X, Y, and Z axes and visualizing 3D FFT distributions to assess frequency content uniformity. |
| | Large-scale slice | 40 | Random non-repeated slices from six public electron tomography datasets. | | |
| | Low-resolution organelle | 40 | Randomly selected non-repeated segmentation results from slices of six public electron tomography datasets. | | |

**Supplementary Table 3 Ablation study of DF5T.** This table examines the contributions of TRformer, Hybridsample, and tp-DMTA to DF5T's performance.

| | PSNR | SSIM | LPIPS |
|---|---|---|---|
| Baseline | 15.06 | 0.53 | 0.47 |
| Baseline<br>+TRformer | 20.92 | 0.57 | 0.33 |
| Baseline<br>+TRformer<br>+Hybridsample | 21.08 | 0.61 | 0.31 |
| Baseline<br>+TRformer<br>+Hybridsample<br>+tp-DMTA | 21.33 | 0.63 | 0.28 |

### Supplementary Methods

### Degradation Methods for Each Tasks

#### Denoising Degradation

To model noise accumulation in low-dose EM imaging, where electron scattering and detector noise obscure organelle details, each patch is blurred with an isotropic 2D Gaussian kernel with standard deviation $\sigma = 1.2$ and then corrupted by additive Gaussian noise. The degradation model is

$$y = Hx + \epsilon, \quad (1)$$

where $H$ is the separable 2D Gaussian convolution operator with $\sigma = 1.2$, and $\epsilon \sim \mathcal{N}(0, \sigma_0^2 I)$ ($\sigma_0$ is the noise scaling factor during sampling). The 1D Gaussian kernel is constructed as

$$g(k) = \exp\left(-\frac{k^2}{2\sigma^2}\right), \quad \sigma = 1.2, \quad (2)$$

with kernel size automatically determined as the standard way (approximately $8\sigma + 1 = 11$, odd). The kernel is normalized by its sum. The full 2D convolution matrix $H$ admits an exact SVD through the Kronecker product of the two identical 1D Toeplitz convolution matrices.

During training and inference, the forward degradation is $y_0 = Hx + \sigma_0 n$, $n \sim \mathcal{N}(0, I)$, followed by pseudoinverse projection $H^+ y_0$ as the diffusion starting point.

#### Super-resolution

To mimic low-resolution EM imaging, where the electron microscope's point spread function (PSF) and detector pixel binning jointly obscure fine organelle details, each high-resolution patch is first blurred with a Gaussian kernel and subsequently downsampled by an integer factor r using pixel averaging.

$$\bar{\boldsymbol{x}} = \boldsymbol{x} * G(\sigma), \tag{3}$$

where $G(\sigma)$ is an isotropic 2D Gaussian kernel with standard deviation $\sigma = 0.8$ pixels (chosen to reflect the moderate optical blurring in typical EM systems), and $*$ denotes 2D convolution.

Downsampling by factor r is then performed via local averaging:

$$\boldsymbol{y} = \downarrow_r (\bar{\boldsymbol{x}}) = A\bar{\boldsymbol{x}}, \tag{4}$$

where $A \in \mathbb{R}^{d_{\mathrm{lr}}^2 \times d^2}$ is the averaging operator that maps each $r \times r$ block of pixels to its mean, and $d_{\mathrm{lr}} = d / r$ is the low-resolution spatial dimension.

The composite forward degradation operator is therefore

$$H = A \cdot (\cdot * G(\sigma)), \tag{5}$$

which admits an exact SVD representation through separable construction. Specifically, the 1D averaging kernel $\mathbf{h} = \frac{1}{r}\mathbf{1}_{r\times 1}$ is used to build Toeplitz matrices for downsampling, while the Gaussian kernel is similarly decomposed separably. The singular values $\boldsymbol{\sigma}$ are obtained from the Kronecker product of the singular values of the respective 1D operators and repeated across channels.

Finally, additive Gaussian noise models residual detector noise:

$$\boldsymbol{y} = H(\boldsymbol{x}) + \sigma_0 \mathcal{N}(\boldsymbol{0}, I) \tag{6}$$

During training and inference, restored patches are upsampled bilinearly by factor r before stitching to reconstruct the full field of view.

### Deblurring degradation

For the deblurring task we explicitly model blur induced by the contrast transfer function (CTF) of the microscope optics. Along a 1D spatial frequency axis $k$ (in \AA $^{-1}$), given electron wavelength $\lambda$, defocus $\Delta f$, spherical aberration $C_s$, amplitude contrast $a$, phase shift $\phi$, and B-factor $B$, the phase function is[5]:

$$\chi(k) = \pi\lambda\Delta f\, k^2 - \frac{\pi}{2} C_s \lambda^3 k^4 + \phi., \tag{7}$$

The corresponding CTF is

$$\mathrm{CTF}(k) = -\left(\sqrt{1-a^2}\sin\chi(k) + a\cos\chi(k)\right)\exp\left(-\frac{Bk^2}{4}\right)., \tag{8}$$

This oscillatory transfer function captures the periodic inversion and attenuation of membrane contrast as a function of frequency, rather than treating blur as a simple low-pass Gaussian filter. We transform the CTF back to real space to obtain a 1D point spread function(PSF):

$$h_{1\mathrm{D}}(x) = \mathcal{F}^{-1}\{\mathrm{CTF}(k)\}(x), \tag{9}$$

then apply an FFT-shift, center-crop to a kernel of length $K$, and normalize:

$$\sum_x |h_{1\mathrm{D}}(x)| = 1 \tag{10}$$

And a separable 2D PSF is then defined as.

$$H_{\mathrm{CTF}}(x, y) = h_{1\mathrm{D}}(x) h_{1\mathrm{D}}(y), \tag{11}$$

with the associated 2D convolution operator $H_{\text{deblur}}$. Let $H_{\text{small}}$ denote the 1D Toeplitz convolution matrix for $h_{1\text{D}}$, with SVD.

$$. H_{\text{small}} = U_s \operatorname{diag}(s_s) V_s^{\top} . \quad (12)$$

The 2D singular values are then given by the Kronecker product.

$$s_{ij} = s_s^{(i)} s_s^{(j)}, \quad (13)$$

which we flatten and sort to obtain $\mathbf{s}_{\text{deblur}}$. The full deblurring operator is

$$H_{\text{deblur}} = U \operatorname{diag}(\boldsymbol{s}_{\text{deblur}}) V^{\top} . \quad (14)$$

The forward model is

$$\boldsymbol{y}_{\text{deblur}} = H_{\text{deblur}} \boldsymbol{x} + \sigma_0 \epsilon, \quad \epsilon \sim \mathcal{N}(\boldsymbol{0}, I), \quad (15)$$

In the data-consistency step we employ a Wiener-style Tikhonov pseudo-inverse:

$$H_{\text{deblur}}^{+} \boldsymbol{y} = V \operatorname{diag}\left( \frac{s_k}{s_k^2 + \alpha \sigma_0^2} \right) U^{\top} \boldsymbol{y} \quad (16)$$

where $s_k$ are the elements of $\mathbf{s}_{\text{deblur}}$ and $\alpha > 0$ controls the regularization strength. This explicitly suppresses frequencies at or near CTF zeros, where optical information about membranes and organelle boundaries is fundamentally unrecoverable, and prevents the model from hallucinating arbitrary high-frequency structure in those directions.

### 2D Inpainting degradation

To simulate missing regions in electron microscopy (EM) images, such as those caused by tears in ultrathin sections or debris occluding organelle structures (e.g., mitochondrial membranes or endoplasmic reticulum tubules), the inpainting degradation method processes each image patch to identify and mask biologically plausible membrane discontinuities.

Contrast is enhanced using adaptive histogram equalization (CLAHE, clipLimit=4.0, tileGridSize=(8,8)) to highlight organelle boundaries. Edge detection is performed with Canny (thresholds 10 and 250), followed by morphological closure (3×3 elliptical kernel, 2 iterations) and adaptive Gaussian thresholding (blockSize=11, C=2).

Connected components are computed, and regions with area $\in [1,100]$ pixels and both dimensions <80% of patch size are retained as "tears/debris". The largest such regions are eroded (1 iteration) to form the final binary membrane mask $M$. The degradation matrix models mild blurring followed by masking:

$$y = M \odot (K * x) + \epsilon , \tag{17}$$

where $K$ is an isotropic 2D Gaussian kernel with $\sigma = 1.0$ and kernel size $k = 3$:

$$K(x, y) = \exp\left(-\frac{x^2 + y^2}{2 \cdot 1.0^2}\right), \tag{18}$$

normalized by sum. The 1D kernels are

$$g_x(i) = g_y(i) = \exp\left(-\frac{i^2}{2 \cdot 1.0^2}\right), \tag{19}$$

(padded to size 3 and normalized). Toeplitz matrices $H_x$, $H_y$ are built separably,

SVD computed, and overall singular values are the outer product $S_x \otimes S_y$, repeated across channels. Gaussian noise $\epsilon \sim \mathcal{N}(0, \sigma_0^2 I)$ is added after masking.

**3D Isotropic degradation**

To emulate anisotropic resolution along the $z$ axis (e.g., due to thicker sections or limited tilt range), we start from a 3D anisotropic Gaussian kernel

$$G_{3\mathrm{D}}(x, y, z) = \frac{1}{Z_{xyz}} \exp\left( -\frac{x^2}{2\sigma_x^2} - \frac{y^2}{2\sigma_y^2} - \frac{z^2}{2\sigma_z^2} \right), \tag{20}$$

with $\sigma_z > \sigma_x, \sigma_y$ to reflect poorer resolution along $z$. Discretely, $G_{3\mathrm{D}}$ is defined on a $K \times K \times K$ grid and normalized6. We then sum over \$z\$ to obtain an effective 2D kernel:

$$K_{\mathrm{iso}}(x, y) = \sum_z G_{3\mathrm{D}}(x, y, z), \qquad K_{\mathrm{iso}}(x, y) \leftarrow \frac{K_{\mathrm{iso}}(x, y)}{\sum_{x,y} K_{\mathrm{iso}}(x, y)}, \tag{21}$$

Let $H_{\mathrm{iso}}$ denote the associated 2D convolution operator, with singular values obtained as in previous tasks via the SVD of the 1D Toeplitz matrices along $x$ and $y$ (denoted $H_x$ and $H_y$ if needed).

For cryo-electron tomography, a single tilt series corresponds to one 3D volume rather than a temporal sequence. To avoid $z$-direction data leakage, we treat each slice $x^{(t)}$ independently and do \emph{not} use any cross-slice information in the forward model:

$$\boldsymbol{y}_{\mathrm{iso}}^{(t)} = H_{\mathrm{iso}} \boldsymbol{x}^{(t)} + \sigma_0 \boldsymbol{\epsilon}^{(t)}, \quad \boldsymbol{\epsilon}^{(t)} \sim \mathcal{N}(\boldsymbol{0}, I), \tag{22}$$

the pseudo-inverse used in data consistency is:

$$H_{\mathrm{iso}}^{+}\boldsymbol{y}^{(t)} = V\,\mathrm{diag}\left(\frac{1}{s_k+\lambda}\right)U^{\top}\boldsymbol{y}^{(t)}. \tag{23}$$

Crucially, we only enforce consistency in frequency directions where $s_k > 0$, and we do not attempt to fill missing wedge/cone regions in Fourier space, where the tomogram contains no physical information.

For serial block-face volume EM (FIB-SEM or SBEM), adjacent slices correspond to physically distinct sections of the specimen (e.g., successive layers of mitochondria or ER). In this case, it is physically meaningful to exploit cross-slice context. If explicitly enabled, we construct an intermediate slice

$$\tilde{x}^{(t)} = (1-\lambda^{(t)})x^{(t)} + \lambda^{(t)}x^{(t-1)} + \gamma\Phi\left(x^{(t)}, x^{(t-1)}\right) \tag{24}$$

where $\lambda^{(t)} = \lambda_0 \mathrm{SSIM}\left(x^{(t)}, x^{(t-1)}\right)$ is an SSIM-based adaptive weight, and $\Phi(\cdot,\cdot)$ is a VGG16 feature similarity term with weight $\gamma$. The forward model then becomes

$$\boldsymbol{y}_{\mathrm{iso}}^{(t)} = H_{\mathrm{iso}}\,\boldsymbol{x}^{(t)} + \sigma_0\epsilon^{(t)} \tag{25}$$

This design encourages consistency of membranes and organelles across consecutive vEM slices, while still respecting the physical acquisition model. For cryo-ET, we do not enable this cross-slice fusion.

### Performance analysis

For the five tasks, we evaluated performance using distinct metrics tailored to each dataset, with detailed results presented in Table 2.

PSNR (Peak Signal-to-Noise Ratio) measures restoration accuracy; higher values indicate a restored image closer to the ground truth (GT):

$$\mathrm{PSNR} = 10\log_{10}\left(\frac{L}{\mathrm{MSE}}\right), \tag{26}$$

where $\mathrm{MSE} = \frac{1}{N}\sum_{i=1}^{N}\left(y(i) - \hat{y}(i)\right)^2$, $y(i)$ is the $i$-th pixel in $y$, and $L$ is the maximum pixel value.

SSIM (Structure Similarity Index Measure) evaluates structural similarity between a restored image $\hat{y}$ and GT image $\mathrm{y}$:

$$\mu_{\hat{y}} = \frac{1}{N}\sum_{i=1}^{N}\hat{y}(i), \tag{27}$$

$$\sigma_y = \sqrt{\frac{1}{N-1}\sum_{i=1}^{N}\left(\hat{y}(i) - \mu_{\hat{y}}\right)^2}, \tag{28}$$

$$\sigma_{y\hat{y}} = \frac{1}{N-1}\sum_{i=1}^{N}\left(y(i) - \mu_y\right)\left(\hat{y}(i) - \mu_{\hat{y}}\right), \tag{29}$$

$$C_l(y,\hat{y}) = \frac{2\mu_y\mu_{\hat{y}} + (k_1L)^2}{\mu_y^2 + \mu_{\hat{y}}^2 + (k_1L)^2}, \tag{30}$$

$$C_c(y,\hat{y}) = \frac{2\sigma_y\sigma_{\hat{y}} + (k_2L)^2}{\sigma_y^2 + \sigma_{\hat{y}}^2 + (k_2L)^2}, \tag{31}$$

$$C_s(y,\hat{y}) = \frac{\sigma_{y\hat{y}} + C}{\sigma_y\sigma_{\hat{y}} + C}, \tag{32}$$

$$\mathrm{SSIM}(y,\hat{y}) = C_l(y,\hat{y})C_c(y,\hat{y})C_s(y,\hat{y}), \tag{33}$$

where $C_l$, $C_c$, and $C_s$ represent brightness, contrast, and structure comparisons. Constants $k_1 = 0.01$, $k_2 = 0.03$, and $C = \frac{(k_2 L)^2}{2}$ ensure stability. SSIM ranges from 0 to 1, with higher values indicating better similarity.

LPIPS (Learned Perceptual Image Patch Similarity) measures the perceptual similarity between a reference image $x$ and a target image $x_0$ using features extracted from a pre-trained deep neural network. A lower LPIPS value indicates greater perceptual similarity:

$$d(x, x_0) = \sum_l \frac{1}{H_l W_l} \sum_{h,w} \| w_l \odot \left( \hat{y}^l_{hw} - \hat{y}^l_{0hw} \right) \|_2^2, \quad (34)$$

where $\frac{1}{H_l W_l}$ denotes the spatial normalization coefficient, $\sum_{h,w}$ represents the summation over spatial positions(h,w)，$\hat{y}^l_{hw}$ and $\hat{y}^l_{0hw}$ are the feature representations being compared. The metric integrates differences across multiple levels (edges, textures, and semantics), with learned weights automatically emphasizing salient features.

Denoising：Normalized Membrane Gradient Profile Analysis. To evaluate edge strength and clarity along a designated membrane boundary, a gradient-based analysis was performed. The gradient magnitude is computed to quantify edge sharpness, with normalization applied to enable comparison across methods. A lower normalized gradient magnitude with aligned peak positions indicates superior edge-preserving performance:

$$\begin{cases} G_x = \text{Sobel}_x(I, \text{ksize} = 3) \\ G_y = \text{Sobel}_y(I, \text{ksize} = 3) \end{cases} \quad (35)$$

$$M = \sqrt{G_x^2 + G_y^2} \quad (36)$$

$$R = M[y_1 - 5 : y_1 + 5, x_1 : x_2] \quad (37)$$

$$P = \frac{1}{10} \sum_{i=-5}^{4} R[i,:] \quad (38)$$

$$P_{\text{norm}} = \frac{P - \min(P)}{\max(P) - \min(P) + \epsilon} \quad (39)$$

$$(x_s, P_s) = \text{interp1d}(x, P_{\text{norm}}, \text{kind}) \quad (40)$$

where I is the input grayscale image, $G_x$ and .$G_y$. are the gradients in the x- and y-directions computed using a 3x3 Sobel filter, M is the gradient magnitude, R is the region of interest (ROI) extracted along the boundary from coordinates $(x_1, y_1)$ to $(x_2, y_2)$ with a 10-pixel vertical span, P is the average gradient magnitude profile across the 10-pixel heigh, $P_{\text{norm}}$ is the min-max normalized profile with $\epsilon = 10^{-6}$ to prevent division by zero, and $(x_s, P_s)$ represents the smoothed profile obtained via cubic interpolation over 200 evenly spaced points.

Fourier spectrum analysis was utilized to assess the frequency distribution of images by computing the Fourier transform and analyzing the intensity of the resulting spectrum to quantify image details and noise levels. The spectrum mean correlates with the preservation of high-frequency details:

$$F = \mathrm{FFT2}(I)\,, \quad (41)$$

$$F_{\mathrm{shift}} = \mathrm{FFTShift}(F)\,, \quad (42)$$

$$S = \log(|F_{\mathrm{shift}}| + 1)\,, \quad (43)$$

$$M_f = |F_s|\,, \quad (44)$$

$$\text{HF Ratio} = \frac{\sum_{(x,y)\in\Omega} M_f(x,y)}{\sum_{(x,y)} M_f(x,y)}\,, \quad (45)$$

where I is the input grayscale image, F is the 2D Fourier transform of I, $F_s$ is the shifted Fourier spectrum with zero frequency at the center, $M_f$ is the magnitude of the Fourier spectrum, S is the logarithmic transformation of the magnitude for visualization, and $\text{HF Ratio}$ is the ratio of the sum of $M_f$ over the high-frequency region $\Omega$ (defined as pixels outside a circular region with radius $\min(h,w)/4$, where h and w are the image height and width) to the total sum of $M_f$. The region $\Omega$ is defined by the mask $(x-c_x)^2 + (y-c_y)^2 > (\min(h,w)/4)^2$, where $(c_x, c_y) = (w/2, h/2)$.

Super-resolution: FRC measures the spatial frequency correlation between a reference image $I_{\mathrm{ref}}$ and a target image $I_{\mathrm{target}}$, often used to estimate resolution in image processing. A higher FRC value at a given frequency indicates better similarity:

$$\mathrm{I}_{\mathrm{ref}}^{\mathrm{ring}} = \frac{\mathrm{I}_{\mathrm{ref}}^{amp}[\mathrm{r}] - \mu_{\mathrm{ref}}}{\sigma_{\mathrm{ref}} + \epsilon}\,, \quad (46)$$

$$I^{\mathrm{ring}}_{\mathrm{target}} = \frac{I^{amp}_{\mathrm{target}}[\mathrm{r}]\text{-}\mu_{\mathrm{target}}}{\sigma_{\mathrm{target}} + \epsilon}, \quad (47)$$

$$\mathrm{FRC}(r) = \frac{\sum I^{\mathrm{ring}}_{\mathrm{ref}} \cdot I^{\mathrm{ring}}_{\mathrm{target}}}{\sqrt{\sum I^{\mathrm{ring}2}_{\mathrm{ref}} \cdot \sum I^{\mathrm{ring}\ 2}_{\mathrm{target}}}}, \quad (48)$$

$$I^{amp}_{ref} = |\mathrm{shift}(\mathrm{FFT2}(I_{\mathrm{ref}}))| \ , \quad (49)$$

$$I^{amp}_{\mathrm{target}} = |\mathrm{shift}(\mathrm{FFT2}(I_{\mathrm{target}}))|, \quad (50)$$

$$\mathrm{Resolution} = \mathrm{interp}(\mathrm{FRC}, 0.143), \quad (51)$$

where $I_{\mathrm{ref}}$ and $I_{\mathrm{target}}$ are the reference and target images, $\mathrm{FFT2}$ computes the 2D Fourier Transform, $\mathrm{FFTShift}$ shifts the zero-frequency component to the center, $r$ 、 $\mu$ and $\sigma$ are the mean and standard deviation of the amplitudes in the ring, $\sigma$ is a small constant (e.g., $10^{-6}$) to avoid division by zero, $\mathrm{f(r)}$ converts radius to spatial frequency, and $\mathrm{interp}$ is determined by linear interpolation at a threshold (e.g., 0.143). In the formula, $\mathrm{FRC}$ employs a normalized amplitude $\tilde{A}$， whereas traditional FRC directly uses the Fourier amplitude. This normalization mitigates the impact of noise and intensity variations, thereby enhancing the accuracy of FRC in evaluating true structural resolution.

Deblurring: Edge Sharpness quantifies the clarity of edges in an image by analyzing the gradient magnitudes at detected edge locations. Higher sharpness values indicate clearer edges:

$$I_{\mathrm{norm}} = \mathrm{Normalize}(I, [0, 255]) \quad (52)$$

$$E = \text{Canny}(I_{\text{norm}}, \sigma = 1) \tag{53}$$

$$G = \text{Sobel}(I_{\text{norm}}) \tag{54}$$

$$G_E = \{G(x,y) | \; E(x,y)\} \tag{55}$$

$$G_{\text{trimmed}} = \text{sort}(G_E)[0.1 \cdot | \; G_E | : 0.9 \cdot | \; G_E |] \tag{56}$$

$$G_{\text{trimmed}} = \text{clip}(G_{\text{trimmed}}, 0, \text{percentile}(G, 95)) \tag{57}$$

$$\text{Edge Sharpness} = \text{mean}(G_{\text{trimmed}}), \text{ if } |G_E| > 0, \text{else} 0 \tag{58}$$

$$\text{Edge SharpnessP95} = \text{percentile}(G_{\text{trimmed}}, 95), \text{ if } |G_E| > 0, \text{else} 0 \tag{59}$$

where I is the input grayscale image, $I_{\text{norm}}$ is normalized to [0, 255], E is the Canny edge map, G is the Sobel gradient magnitude, $G_E$ are gradients at edges, $G_{\text{trimmed}}$ is the trimmed gradient array, and $\text{Edge Sharpness Median}_{\text{orig}}$ is the original image's median sharpness (1 if zero).

2D Inpainting: Membrane Sharpness. Quantifies edge sharpness by computing the average gradient magnitude across the image.

$$G_x = \text{Sobel}_x(I, \text{ksize} = 3), \tag{60}$$

$$G_y = \text{Sobel}_y(I, \text{ksize} = 3), \tag{61}$$

$$M = \sqrt{G_x^2 + G_y^2}, \tag{62}$$

$$\text{Membrane Sharpness} = \frac{1}{N}\sum_{(x,y)} M(x,y), \tag{63}$$

where I is the input grayscale image, $G_x$ and $G_y$ are the gradients in the x- and y-directions computed using a 3x3 Sobel filter, M is the gradient magnitude, and Membrane Sharpness is the mean of M over all N pixels in the image.

Organelle Contrast: Measures the intensity difference between foreground (organelles) and background regions, segmented using Otsu's thresholding.

$$T = \text{OtsuThreshold}(I) \tag{64}$$

$$F = \{I(x,y) | \; I(x,y) \geq T\} \tag{65}$$

$$B = \{I(x,y) | \; I(x,y) < T\} \tag{66}$$

$$\text{OrganelleContrast} = \frac{1}{|F|}\sum_{I(x,y)\in F} I(x,y) - \frac{1}{|B|}\sum_{I(x,y)\in B} I(x,y) \tag{67}$$

where T is the threshold determined by Otsu's method, F is the set of foreground pixels ($I(x,y) \geq T$), B is the set of background pixels ($I(x,y) < T$), and Organelle Contrast is the difference between the mean intensities of the foreground and background regions.

Structural Integrity: Evaluates the structural similarity between the reference and test images by comparing segmented regions obtained via watershed segmentation.

$$M_{r,t} = \begin{cases} 1 & \text{if } I_{r,t}(x,y) < 30, \\ 2 & \text{if } I_{r,t}(x,y) > 200, \\ 0 & \text{otherwise,} \end{cases} \tag{68}$$

$$S_{r,t} = \text{Watershed}(I_{r,t}, M_{r,t}) \tag{69}$$

$$O = \sum_{(x,y)} 1\{S_r(x,y) > 1 \wedge S_t(x,y) > 1\} \tag{70}$$

$$U = \sum_{(x,y)} 1\{S_r(x,y) > 1 \vee S_t(x,y) > 1\} \tag{71}$$

$$\text{Structural Integrity} = \begin{cases} \dfrac{O}{U} & \text{if } U > 0, \\ 0 & \text{otherwise,} \end{cases} \tag{72}$$

where $I_r$ and $I_t$ are the reference and test grayscale images, $M_r$ and $M_t$ are marker arrays for watershed segmentation based on intensity thresholds, $S_r$ and $S_t$ are the segmented regions, O is the number of overlapping foreground pixels ($S_r > 1$ and $S_t > 1$), U is the number of pixels in the union of foreground regions, and $\text{Structural Integrity}$ is the intersection-over-union (IoU) of the segmented foreground regions.

Edge Preservation: Measures the proportion of edges in the reference image that are retained in the deblurred image using Canny edge detection.

$$E_{r,t} = \text{Canny}(I_{r,t}, \sigma = 1) \tag{73}$$

$$\text{Edge Preservation} = \frac{\sum_{(x,y)} 1\{E_r(x,y) \wedge E_t(x,y)\}}{\sum_{(x,y)} 1\{E_r(x,y)\}} \tag{74}$$

where $E_r$ and $E_t$ are binary edge maps of the reference and test images, respectively, computed using Canny edge detection with $\sigma = 1$, and $\text{Edge Preservation}$ is the ratio of overlapping edge pixels to the total number of edge pixels in $E_r$.

3D Isotropic: To evaluate the effectiveness of Z-axis anisotropy restoration in volume electron microscopy, we conducted a detailed Fourier Ring Correlation (FRC) analysis on the YZ and ZX planes. By comparing the FRC curves and the amplitude distributions of the 3D Fourier transforms between the original and restored data, we quantified the impact of the anisotropy restoration:

$$FRC(r) = \frac{\sum_{k \in \text{ring}(r)} \tilde{F}_1(k) \cdot \tilde{F}_2(k)}{\sqrt{\sum_{k \in \text{ring}(r)} \tilde{F}_1(k)^2} \cdot \sqrt{\sum_{k \in \text{ring}(r)} \tilde{F}_2(k)^2}}, \quad (75)$$

where $\tilde{F}_1(k) = \frac{|F_1(k)| - \mu_1}{\sigma_1 + \epsilon}$, $\tilde{F}_2(k) = \frac{|F_2(k)| - \mu_2}{\sigma_2 + \epsilon}$, $|F_1(k)|$ and $|F_2(k)|$ are the magnitudes of the Fourier transforms of the reference and target images, $\mu_1$, $\mu_2$ are the means of the magnitudes in the ring, $\sigma_1$, $\sigma_2$ are the standard deviations of the magnitudes in the ring, $\epsilon = 10^{-6}$ is a small constant to avoid division by zero.

**Supplementary References**


1. Lu, C. *et al.* Diffusion-based deep learning method for augmenting ultrastructural imaging and volume electron microscopy. *Nat. Commun.* **15**, 4677 (2024).
2. Xia, J. *et al.* Blind Super-Resolution via Meta-Learning and Markov Chain Monte Carlo Simulation. *IEEE Trans. Pattern Anal. Mach. Intell.* **46**, 8139–8156 (2024).
3. Ma, C., Tan, W., He, R. & Yan, B. Pretraining a foundation model for generalizable fluorescence microscopy-based image restoration. *Nat. Methods* **21**, 1558–1567 (2024).
4. Stringer, C. & Pachitariu, M. Cellpose3: one-click image restoration for improved cellular segmentation. *Nat. Methods* **22**, 592–599 (2025).
5. Guo, M. *et al.* Deep learning-based aberration compensation improves contrast and resolution in fluorescence microscopy. *Nat. Commun.* **16**, 313 (2025).
6. Popovych, S. *et al.* Petascale pipeline for precise alignment of images from serial section electron microscopy. *Nat. Commun.* **15**, 289 (2024).
